# Genome-wide analysis of phased small interfering RNAs related to tomato fruit ripening and quality

**DOI:** 10.1101/2025.03.20.644405

**Authors:** Santiago Stalder, Agustín Sabbione, Lucas Daurelio, Marcela Dotto

## Abstract

Fruit quality and sensorial aspects are key determinants for costumer selection and satisfaction. Main characteristics include texture, color, aroma and taste, which are defined by a wide range of molecular processes that will ultimately determine fruit general appearance. The production of cultivars with optimized metabolite contents is increasingly being explored, in particular through the study of gene regulatory networks that modulate the metabolism of these compounds. There are several types of small RNAs originating from different biosynthesis mechanisms that regulate a large number of genes involved in various processes during plant development and growth. However, reports on their participation in regulatory networks related to qualitative aspects of fruits are scarce. In this work, genome-wide bioinformatics analyses were performed to identify regulatory modules that involve the action of phased secondary small interfering RNAs in different ripening stages of tomato fruits. We identified small RNA regulatory modules related to the acquisition of commercially important traits, such as the development of brightness, color, firmness, and nutritional content, which set the basis for further characterization of phasiRNA-mediated regulation of aspects related to tomato fruit quality improvement and the possibility of using these regulatory RNAs as tools for molecular breeding.

## 1. Introduction

Tomato (*Solanum lycopersicum* L.) is one of the main vegetable crops produced and consumed worldwide. During the ripening process fruits develop and undergo a series of physiological changes that make them attractive to their consumers: they grow in size, slowly soften, change their colors, produce characteristic aromas and acquire pleasant flavors. These changes contribute to the development of organoleptic properties that need to be optimized for consumer’s acceptance. Properties such as good taste, pleasant aroma, intense red color and firm texture are amongst consumeŕs preferences for fresh tomato fruits (Baldwin *et al*. 1998; Causse *et al*. 2010). All these aspects are determined by a complex combination of metabolites, mainly combination of carbohydrates, organic acids, amino acids and a number of volatile compounds, mostly derived from phenylpropanoids (Tieman *et al*. 2012; Peled-Zehavi *et al*. 2015).

In addition, hormones are also important factors contributing to ripening and to the development of organoleptic properties. Being a climacteric fruit, ethylene is key during ripening, contributing to cell wall disassembly leading to changes in fruit texture and firmness, as well as changes in volatile compounds and soluble solids, promoting delightful aroma and flavor (Alexander 2002). In addition, auxin, ABA and jasmonic acid also influence expression of genes involved in the biosynthesis of ethylene and other aspects of the ripening control network (Li *et al*. 2021).

Changes underlying the dynamic metabolism controlling these aspects are in most cases related to the regulation of gene expression of key steps of metabolic pathways. Therefore, regulation of gene expression has been extensively studied in tomato fruit ripening, including several studies to investigate the role of microRNAs (miRNAs) and other small regulatory RNAs (Klee and Giovannoni 2011; Mohorianu *et al*. 2011; Zuo *et al*. 2012; Karlova *et al*. 2013; Din and Barozai 2014; Azzi *et al*. 2015).

Small regulatory RNA molecules are 20 to 24-nt-long small RNAs that can regulate gene expression at the transcriptional or post-transcriptional level. Most extensively studied are miRNAs, which bind target transcripts through perfect or near-perfect base complementarity as part of an RNA-Induced silencing Complex (RISC) which cleaves and down-regulates target transcript expression. Several miRNAs have been shown to participate in fruit ripening process, mainly through the post-transcriptional regulation through cleavage of transcripts coding for transcription factors (Karlova *et al*. 2013; Din and Barozai 2014). Is important to mention that miRNAs are only a very small fraction within the total small RNAs present in a celĺs transcriptome and the function of other classes of small RNAs is typically less characterized.

Other types of small RNAs include trans-acting short interfering RNAs (tasiRNAs) and phased small interfering RNAs (phasiRNAs). These are 21-nt-long small RNAs which in some cases were also shown to down-regulate target transcripts (Zhang *et al*. 2012; Fei *et al*. 2013a; Yoshikawa 2013; Zheng *et al*. 2014). Previous studies have shown that some phasiRNAs increase their expression in response to pathogen infection, such as *Phytophthora sojae* or *Botrytis cinerea* (Yang and Huang 2014; Wu *et al*. 2017) or in response to drought stress (Sosa-Valencia *et al*. 2016). Moreover, phasiRNA-producing loci were shown to promote flowering and reproduction (Liu *et al*. 2020). In other studies, these small RNAs were shown to be derived from and participate in the regulation of genes coding for NB-LRR (Zhai *et al*. 2011; Fei *et al*. 2013a; Zheng *et al*. 2014) and a particular class of 24-nt-long phasiRNAs were shown to accumulate in reproductive organs in maize, rice and other angiosperms (Zhai *et al*. 2015; Kakrana *et al*. 2018; Xia *et al*. 2019). However, in tomato these reports have mostly been performed using leaf tissue or mature red tomato fruits and at the moment a complete analysis of phasiRNA production throughout tomato fruit ripening is missing. (Wu *et al*. 2017; Luan *et al*. 2020).

Here we conducted genome-wide analyses to identify phased small RNAs during tomato fruit ripening by analyzing data corresponding to four ripening stages: immature green (IG), breaker (Br), pink (Pk) and red ripe (RR) from *S lycopersicum* cv. Ailsa Craig fruits. In addition, we used tomato fruit degradome data to validate target transcripts regulated by phasiRNAs and found phasiRNA/target modules regulating several aspects of fruit growth and development, including genes involved in plant defense, but also in the regulation of fruit ripening and quality, including genes related to fruit brightness, color, texture and acidity. The results presented here set the grounds for further exploration and possibly the development of biotechnological tools to improve organoleptic properties in tomato fruits.

## 2. Materials and methods

### 2.1. Data retrieval

Small RNA sequencing data, RNA-seq and degradome data was downloaded from Sequence Read Archive database (SRA, https://www.ncbi.nlm.nih.gov/sra) using fastq-dump from SRA-toolkit (http://ncbi.github.io/sra-tools/) in order to convert SRA original format into fastq. All sequencing data correspond to samples from tomato fruits (*S. lycopersicum* cv. Ailsa Craig) in different ripening stages (Supplemental File 1). Small RNA-seq analyzed data correspond to two biological replicates from four ripening stages: immature green (IG), breaker (Br), pink (Pk) and red ripe (RR) from Gao et al. (Gao *et al*. 2015). Degradome data correspond to IG and RR fruits, retrieved from Karlova et al. (Karlova *et al*. 2013).

### 2.2. Pre-processing and filtering of fastq files

Quality control of fastq files was assessed using FastQC v0.12.0 (Andrews 2010). Software fastx_clipper v0.0.14 from FASTX-toolkit (Gordon 2011) was used to trim the adapter portion from small RNA-seq reads, using default parameter and specifying options -l 18, -c and - M 5. Next, sequences corresponding to rRNA, tRNA, snRNA and snoRNA were obtained from Rfam 15.0 (Griffiths-Jones 2003) and removed from fastq files using Bowtie v1.3.1 (Langmead 2010), keeping unaligned reads using option --un. Number of reads remaining after each step are summarized in Supplemental File 1.

### 2.3. Identification of *PHAS*

ShortStack 3.8.3 software (Axtell 2013; Johnson *et al*. 2016) was used on fastq files after pre-processing. Mapping was performed using default parameters and specifying the options “-- nohp” and “--ranmax none”. Only *PHAS* loci with DicerCall = 21 were analyzed, with size longer than 100 pb and *P-score* ≥ 25. *PHAS* loci coordinates were defined by comparing phased clusters across all analyzed small RNA-seq libraries and using visualization in IGV viewer. Sequences of phasiRNAs from the identified loci were extracted from ShortStack-generated bam files and named individually for each *PHAS* locus (Supplemental File 2).

### 2.4. phasiRNA quantification and differential expression analysis

DEUS package was used for phasiRNAs quantification which allows using the actual read sequences to determine expression levels (Jeske *et al*. 2019). Filtered count tables to retain those sequences with more than 1 count were used in downstream analysis. Differential expression of phasiRNAs was evaluated using edgeR 3.42.4 (Robinson *et al*. 2010) in RStudio 2023.06.0 software, using R version 4.4.1. Differential expression was analyzed in paired comparisons for the analyzed ripening stages. Significant statistical difference was established for FDR-corrected q-value ≤ 0,05.

### 2.5. Validation of target genes using degradome data and GO term analysis

Target genes for phasiRNAs were validated using degradome data obtained from fruit samples, corresponding to IG and RR stages (Supplemental File 1) and CleaveLand v4.4 software (Addo-quaye *et al*. 2009). Results with p ≤ 0.05 and T-plots in the 0 categories were analyzed and a manual curation was done to select only isolated highest peaks. Go term enrichment analysis was performed using Singular Enrichment Analysis from AgriGO,v2.0 (Tian *et al*. 2017) selecting Transcripts from ITAG4.0 as background. All phasiRNA targets were categorized according to their functional annotation using ITAG 4.0 file from SolGenomics (https://solgenomics.net/).

### 2.6. RNA extraction and cDNA synthesis

Fruits corresponding to the IG and RR stages were collected from tomato plants cv. Ailsa Craig. Total RNA was extracted from pools of 3 fruits pulverized using liquid nitrogen and processed into three biological replicates. Total RNA was isolated from 100 mg of pulverized fruits using TriPure reagent (Roche), followed by RQ1 DNase (Promega) treatment and posterior purification using the PureLink RNA mini kit (Invitrogen).

First strand cDNA was obtained by reverse transcription on 1 μg of DNAse-treated total RNA using ImProm II reverse transcriptase enzyme (Promega). The obtained cDNA was used as a template for quantitative PCR amplification in a AriaMx 1.6 qPCR System (Agilent), using 1 × *PerfectStart* ® Green qPCR SuperMix, (Transgene Biotech). Primers for the genes under study are available upon request. Amplifications were performed under the following conditions: 30 s of denaturation at 94 °C, 40 cycles at 94 °C for 5 s, 50 °C for 15 s, and 72°C for 10 s. Three biological replicates and three technical replicates were performed for each sample. Melting curves were determined by measuring fluorescence with increasing temperature (from 65 to 95 °C). Gene expression levels were normalized to that of tomato *UBIQUITIN3* gene *(UBI3)*. The efficiency of the primers used for qPCR analysis was determined to be close to 100% for all primers used, and therefore, analysis was performed using the 2^−ΔΔCt^ method (Livak and Schmittgen 2001).

## 3. Results

### 3.1. Identification of phased small interfering RNAs during tomato fruit ripening

In order to identify phasiRNA-generating loci during tomato fruit ripening, we analyzed small RNA-seq data from IG, Br, Pk and RR fruits (Suppl. File 1). We identified 25 *PHAS* loci with a *P-score* ≥ 25 and a length above 100 bp (Table 1), in accordance to similar criteria used in previous reports (Feng *et al*. 2019; Nakamura *et al*. 2019; Das *et al*. 2020).

**Table 1:** *PHAS* loci detected in tomato fruits.

| **Locus** | **Genomic coordinates (SL4.0)** | **Associated Gene model** | **Functional annotation** |
| --- | --- | --- | --- |
| *PHAS1* | ch01:4415780-4416701 | Solyc01r001560.1 | lncRNA |
| *PHAS2* | ch02:29160966-29163434 | Solyc02g036270.4.1 | Disease resistance protein |
| *PHAS3* | ch04:373826-376280 | Solyc04g005540.3/<br>Solyc04r018440.1.1 | Pvr4/<br>lncRNA |
| *PHAS4* | ch05:2549891-2552244 | Solyc05g008070.4.1 | Disease resistance protein |
| *PHAS5* | ch05:3872005-3874078 | Solyc05g009630.4<br>(intron) | Disease resistance protein |
| *PHAS6* | ch06:42183189-42183665 |  |  |
| *PHAS7* | ch06:42207523-42208019 | Solyc06r032010.1.1 | lncRNA |
| *PHAS8* | ch06:437853-438581 | Solyc06g005410.3.1<br>(partial overlap) | Unknown protein |
| *PHAS9* | ch06:439060-439697 |  |  |
| *PHAS10* | ch06:704215-704566 | Solyc06r028700.1 | lncRNA |
| *PHAS11* | ch07:4205316-4207077 | Solyc07g009180.1.1 | Disease resistance protein |
| *PHAS12* | ch08:13837317-13838009 |  |  |
| *PHAS13* | ch08:57063495-57067399 | Solyc08r040460.1.1 | lncRNA |
| *PHAS14* | ch08:59208402-59210241 |  |  |
| *PHAS15* | ch09:13657071-13658918 | Solyc09r018220.3 | Tobacco mosaic virus resistance-2 |
| *PHAS16* | ch09:57495810-57497595 | Solyc09g064240.3 | PFKB-like |
|  |  | (intron) | carbohydrate kinase |
| *PHAS17* | ch10:58274198-58274971 | Solyc10g076270.3.1<br>(intron) | Regulatory-associated<br>protein of TOR1 |
| *PHAS18* | ch11:2733015-2740117 | Solyc11g008530.3/<br>Solyc11r051240.1/<br>Solyc11r051250.1 | DCL-2d/<br>lncRNA/<br>lncRNA |
| *PHAS19* | ch11:2747539-2754315 | Solyc11g008540.3.1<br>(introns/exons)/<br>Solyc11r051240.1 | DCL-2b/<br>lncRNA |
| *PHAS20* | ch11:49432234-49436209 | Solyc11g065780.3.1/<br>Solyc11g065790.1/<br>Solyc11g065810.1<br>(+intergenic regions) | CC-NBS-LRR/<br>Disease resistance<br>protein 12/<br>CC-NBS-LRR |
| *PHAS21* | ch11:52640011-52642954 | Solyc11g069990.2<br>(intron/exon) | I2C5 |
| *PHAS22* | ch11:7217527-7218901 | Solyc11g013770.2.1<br>(intron) | Unknown protein |
| *PHAS23* | ch12:2829642-2831621 | Solyc12g009520.2.1 | Receptor-like protein<br>12 |
| *PHAS24* | ch12:641458-644168 | Solyc12g006040.3<br>(+intergenic) | NBS-LRR |
| *PHAS25* | ch12:6447225-6449223 | Solyc12g016220.3.1 | Disease resistance<br>protein |

Among these, 17 *PHAS* loci are located in genomic locations overlapping annotated protein-coding genes (Table 1). Interestingly, ten of these loci overlap genes coding for different proteins involved in the response to pathogen infection, annotates as ‘Disease resistance proteins’, members of NBS-LRR family, TMV resistance-2, Pvr4 and I2C5 proteins (*PHAS2, 3, 4, 5, 11, 15, 20, 21, 24* and *25*). In addition, *PHAS18 and PHAS19* overlap genes coding for sly-DCL2d and sly-DCL2b, respectively and *PHAS8, 16, 17, 22* and *23* overlap different protein-coding annotated genes. Remarkably, *PHAS1, PHAS7, PHAS10* and *PHAS13* seem to be derived from long non-coding RNAs (lncRNAs) annotated in those regions, whereas for some of them lncRNA genes were also annotated overlapping the protein-coding genes (*PHAS3, PHAS18* and *PHAS19*), and the rest of the identified *PHAS* loci are located in genomic regions where no gene annotation exists (*PHAS6, 9, 12* and *4*) (Table 1).

A detailed analysis of the phasiRNAs mapping to these regions indicates that in several cases they are derived from intronic or intergenic regions (*PHAS5, PHAS6, PHAS8, PHAS9, PHAS12, PHAS14, PHAS16, PHAS17, PHAS18, PHAS19, PHAS20, PHAS21, PHAS22* and *PHAS24)*; (Table 1; Supplemental File 3) and in other cases the genomic regions could give rise to overlapping antisense transcripts (*PHAS3, PHAS18, PHAS19)*; (Table 1; Supplemental File 3), suggesting that in those cases, the phasiRNAs could be synthesized by mechanisms different from the canonical process involving the binding of a trigger miRNA to the mRNA transcript for protein-coding genes (Li *et al*. 2012; Xia *et al*. 2015, 2019).

In order to identify whether any of the detected phasiRNAs could be generated by the canonical mechanism, we used degradome data analysis using the full list of tomato miRNAs as queries and the *PHAS* loci sequences as mapping reference for degradome data from fruits (Karlova *et al*. 2013). Using this procedure, we were able to identify that miR482 could be a trigger for the generation of phasiRNAs produced from *PHAS2*, *PHAS14* and *PHAS22* (Fig. 1), but no cleavage event was validated for the rest of the *PHAS* loci.

**Figure 1:**
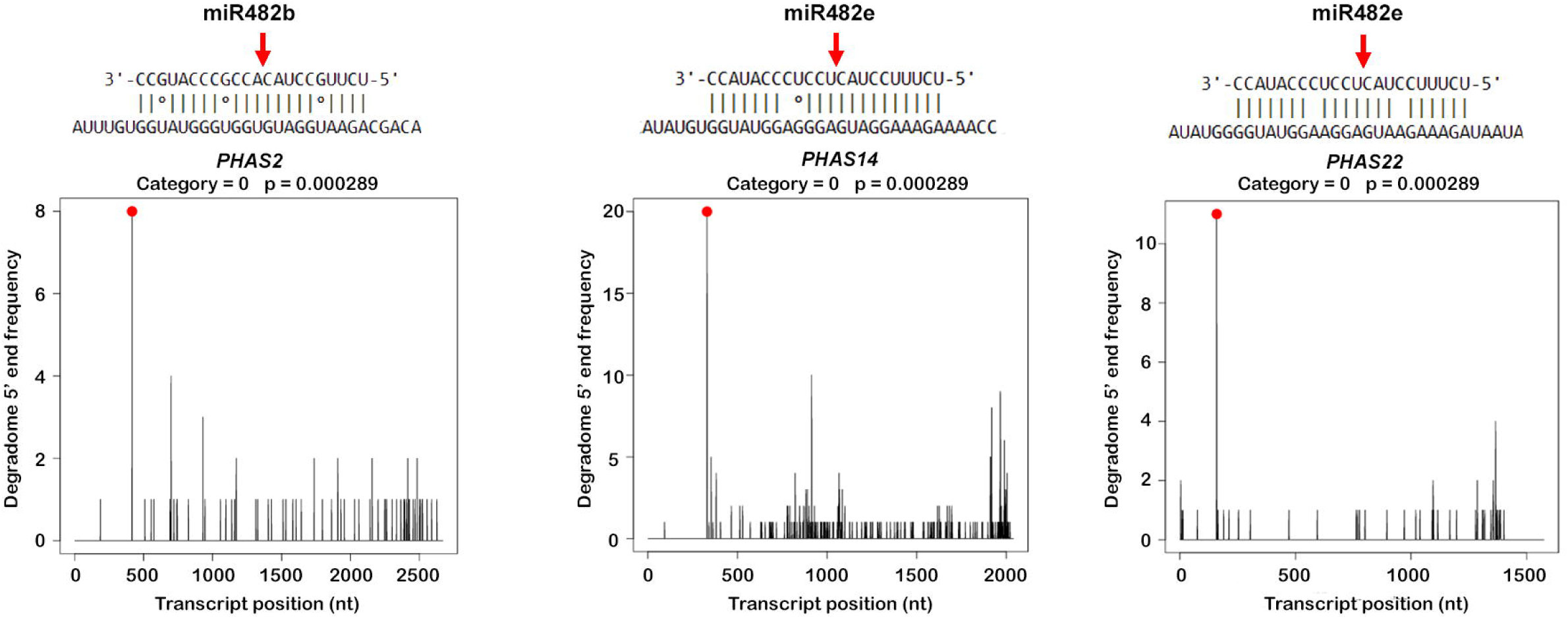
Cleavage validation for *PHAS2, PHAS14* and *PHAS22* transcripts mediated by two members of the miR428 family. Binding sites for miR482b and miR482e on respective *PHAS* transcripts are shown. The red dot in the T-plots indicates normalized reads present in degradome libraries coinciding with predicted cleavage event, signaled with a red arrow on the binding sites. T-plot category and p-value determined by CleaveLand software analysis are indicated.

### 3.2 Prediction and validation of phasiRNA targets

Considering the results from the previous analysis, we performed a genome-wide target prediction and validation of the full-list of phasiRNAs derived from the 25 *PHAS* loci. To do this, we extracted their sequences and named each phasiRNA in relation to the number of *PHAS* loci, start position and DNA strand producing each of them, using the format “Locus_Strand_Order”: L1 to L25 indicates the *PHAS* locus number, the strand is indicated as *S* (sense) or *AS* (antisense) and the order number indicates the phasiRNA start position in the *PHAS* locus (e.g., L1_S_1), (Supplemental File 2).

To investigate the biological importance of these phasiRNAs, we used degradome data analysis to validate target transcripts for all of them, which resulted in a total of 186 validated target transcripts for all phasiRNAs (Supplemental File 4).

Next, we used GO Term Enrichment analysis to identify whether phasiRNAs expressed during tomato fruit ripening might regulate one or a few specific biological processes. Even when there was no significant GO term enrichment, phasiRNAs targets were next classified according to the top levels GO categories ‘Biological Process’, ‘Cellular Component’ and ‘Molecular Function’. We established that the phasiRNAs produced by the 25 identified *PHAS* loci participate in the regulation of several aspects of fruit growth and development, including the terms developmental and metabolic processes; multi-organism and immune system processes (which contemplates interaction with pathogens and defense responses); biological regulation (includes transcription factors), among others (Figure 2).

**Figure 2:**
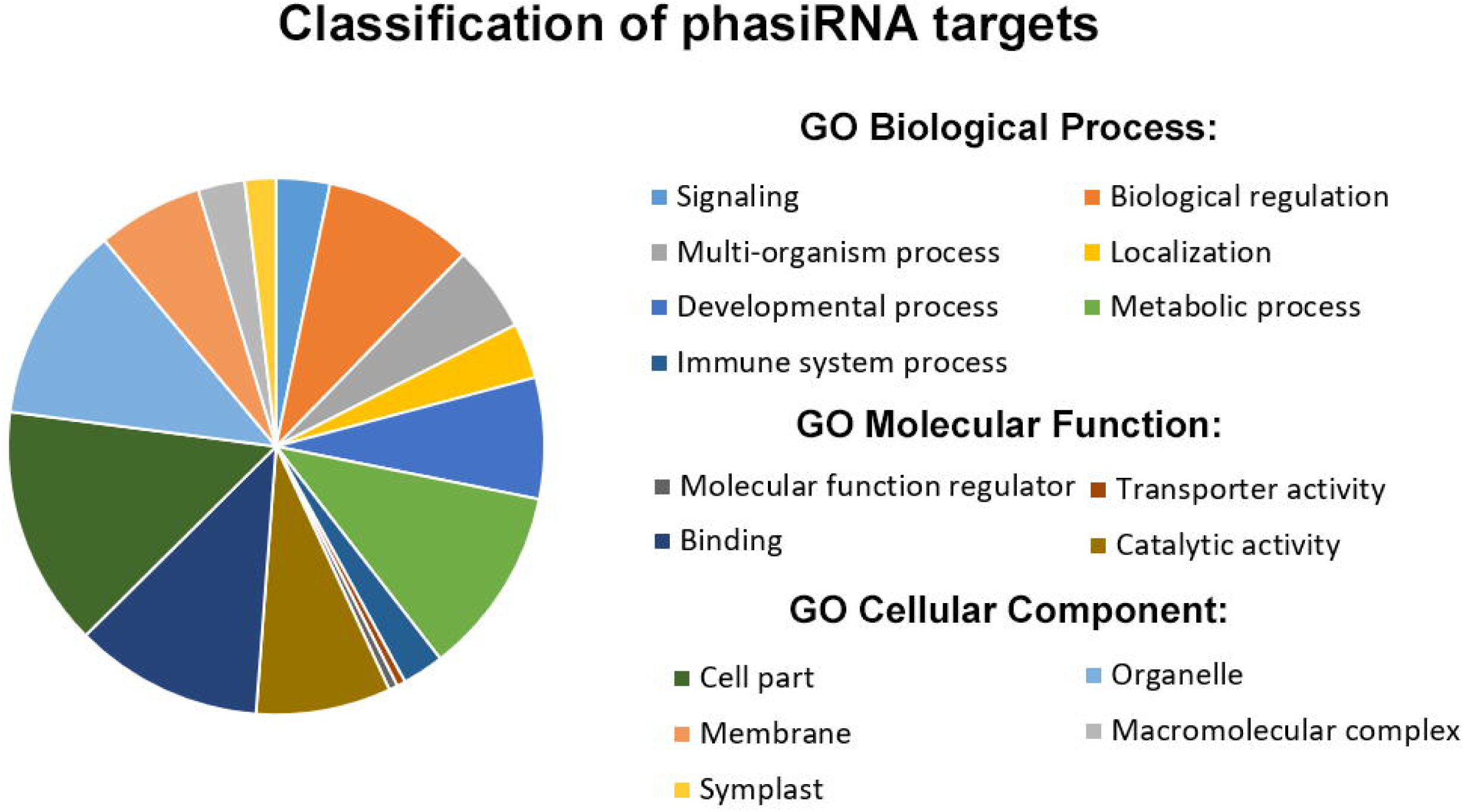
Distribution of the validated target genes for detected phasiRNAs considering top level terms in the GO term annotation categories ‘Biological processes’, ‘Molecular Function’ and ‘Cellular Component’.

Considering one of the most studied aspects of phasiRNAs function is related to their regulation of genes involved in the response to pathogens and plant defense (Fei *et al*. 2013a; Liu *et al*. 2020; Luan *et al*. 2020), we focused our downstream analyses in those targeting genes participating in other aspects of fruit growth and development. In particular, we explored the possibility that some of these phasiRNAs could participate in the regulation of genes involved in fruit ripening and in the processes related to the establishment of fruit quality traits and the development of organoleptic properties. Therefore, we categorized them based on their functional annotations into groups according to their possible role in fruit color and brightness development, acquisition of flavor and aroma, softening and general aspects of fruit ripening, as summarized in Table 2.

**Table 2:**
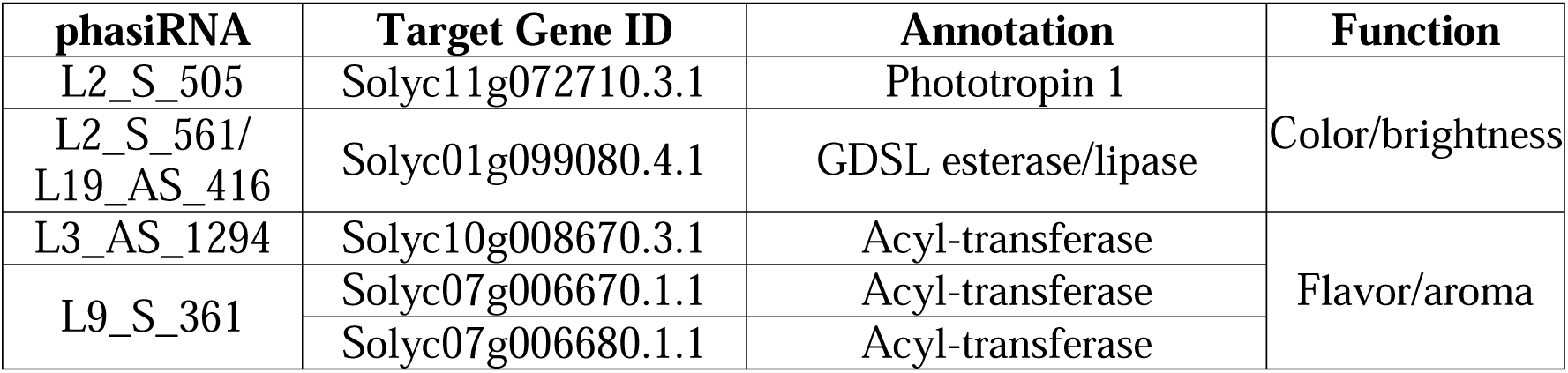
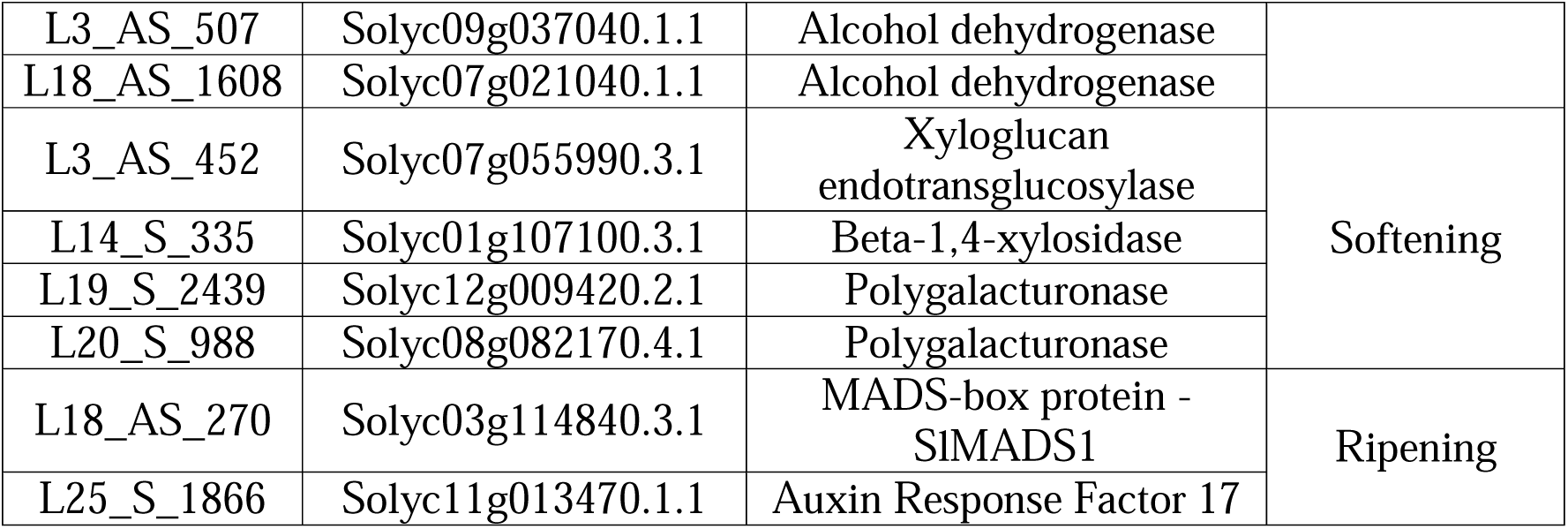
Selected phasiRNAs targeting genes involved in fruit ripening and quality.

#### 3.2.1 PhasiRNAs involved in color and brightness development

Target validation with degradome data analysis allowed the detection of target genes for phasiRNAs derived from *PHAS*2 and *PHAS19* related to the development of fruit color and brightness (Table 2). Target validation for these phasiRNAs indicates that there is a high accumulation of cleaved transcripts at the predicted cleavage sites mediated by these phasiRNAs in the degradome libraries analyzed and this information is shown in T-plots (Fig. 3A-B). PhasiRNA L2_S_505 regulates the transcript coding for Phototropin 1 (Solyc11g072710.3.1; Fig. 3A right), which has been related to carotenoid content and has been proposed as a potential tool for modulation of lycopene content in tomato fruits (Kilambi *et al*. 2021). In addition, two phasiRNAs targeting the same gene coding for a GDSL lipase were validated (Table 2), enzymes that are involved in the formation of the cuticle and brightness of fruits (Yeats *et al*. 2012). These two phasiRNAs, L2_S_561 and L19_AS_416 showed expression levels that remain approximately constant during ripening, increasing slightly the accumulation of L19_AS_416 in the RR stage (Fig. 3B, left). These phasiRNA were validated as regulators of transcript Solyc01g099080.4.1 (Table 2, Fig. 3B, right), with a cleavage site mediated by L2_S_561 in the 3’ region of the transcript (red dot in Fig. 3B) and a cleavage site mediated by L19_AS_416 in the 5’ region of the transcript (blue dot in Fig. 3B).

**Figure 3:**
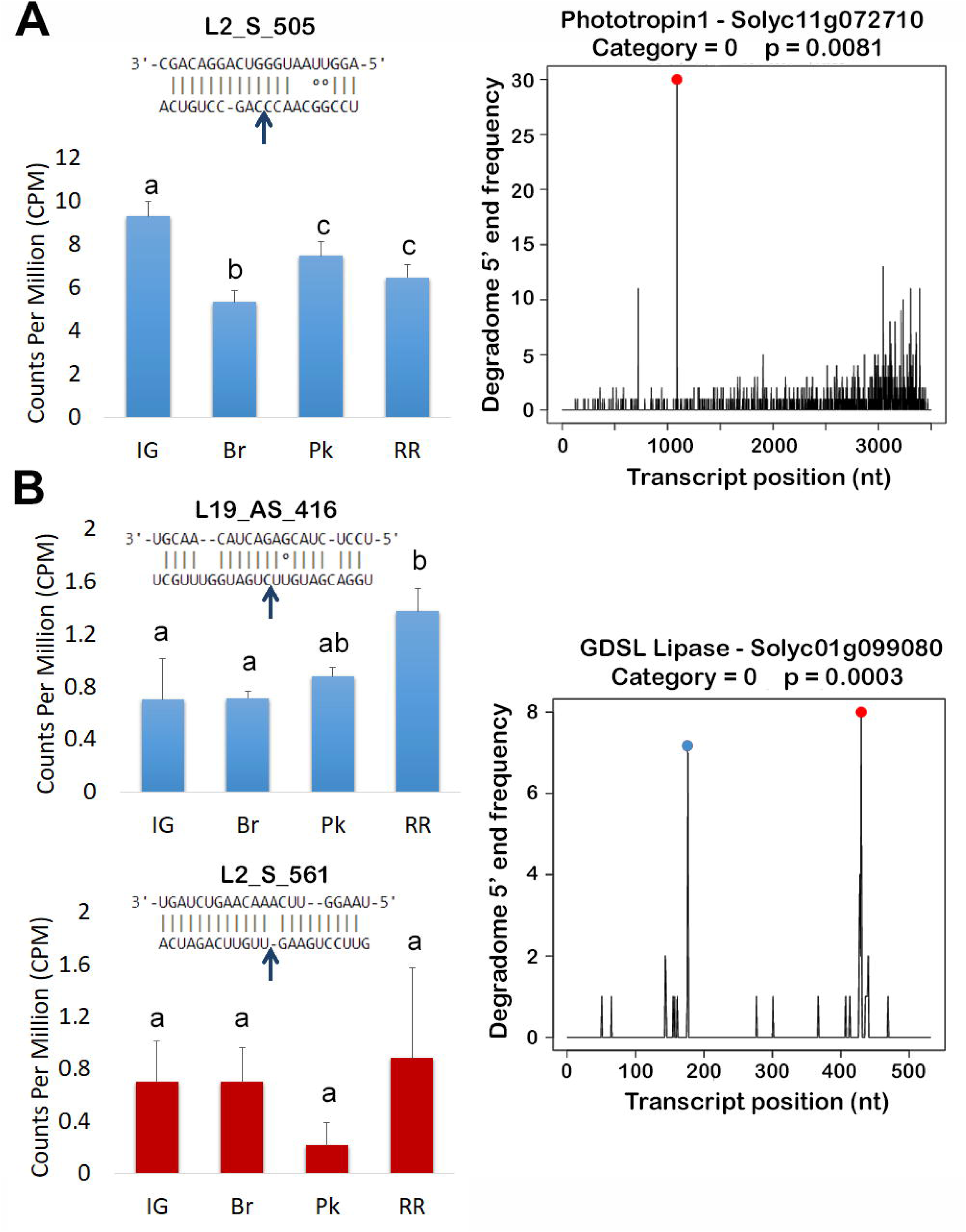
Fruit color and brightness. (A) Expression profile of phasiRNA L2_S_505 in fruits of different ripening stages (left) and validation of the cleavage of transcript Solyc11g072710 coding for Phototropin 1 (right). The red dot indicates the normalized reads that match the predicted cleavage position mediated by L2_S_505 at this transcript in degradome libraries. (B) Expression profile of phasiRNAs L2_S_561 and L19_AS_416 in fruits of different ripening stages (left) and validation of two cleavage events on transcript Solyc01g099080 coding for a GDSL lipase (right). The red and blue dots indicate the normalized reads that match the predicted cleavage positions mediated by L2_S_561 and L19_AS_416, respectively, at this transcript in degradome libraries. Peak category and *p-value* according to CleaveLand are indicated above T-plots. IG: immature green, Br: breaker, Pk: pink, RR: red ripe. Different letters indicate a significant difference (adjusted p-value < 0.05) according to edgeR analysis.

#### 3.2.2 PhasiRNAs involved in fruit flavor and aroma

Additional target transcripts related to the acquisition of fruit flavor and aroma were detected for phasiRNAs derived from locus *PHAS3, PHAS9* and *PHAS18*. L3_AS_507 and L18_AS_1608 regulate the transcripts Solyc09g037040.1.1 and Solyc07g021040.1.1, respectively, coding for Alcohol dehydrogenase (ADH) enzymes and phasiRNAs L3_AS_1294 and L9_S_361 were validated as regulators of genes coding for Acyl transferase (AT) enzymes (Solyc10g008670.3.1, Solyc07g006670 and Solyc07g006690; Table 2).

PhasiRNA L9_S_361 accumulates at lower levels in IG, Br and Pk stages and higher in RR fruits (Fig. 4A, left) and the cleavage point close to the 400^th^ nt in Solyc07g006680.1.1 transcript was validated and is shown as in Fig. 4A. Conversely, L18_AS_1608 exhibits higher expression in the IG stage, decreasing during the rest of the ripening process (Fig. 4B, left), with a validated cleavage point towards the 3’end of the Solyc07g021040.1.1 transcript.

**Figure 4:**
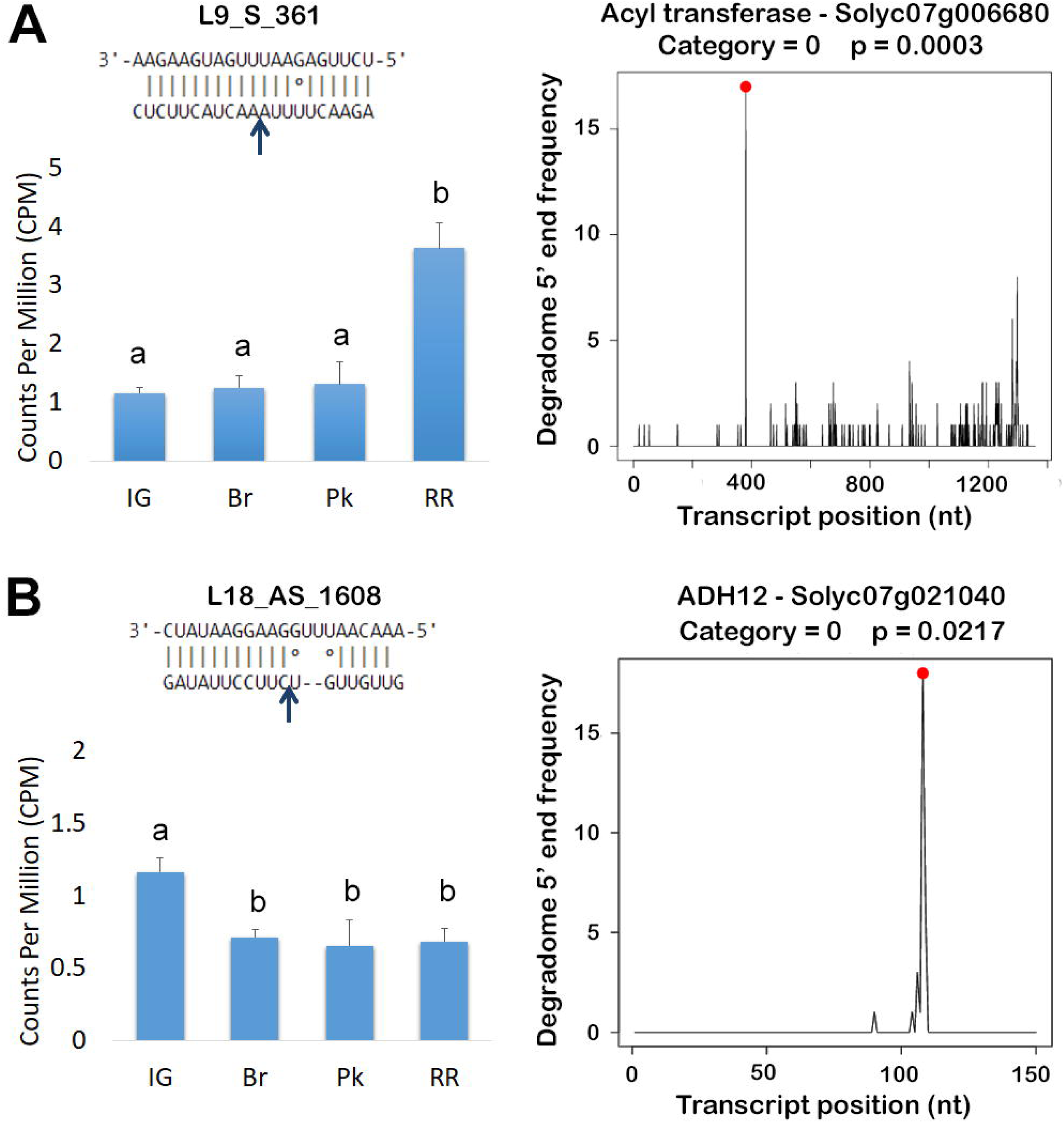
Fruit flavor and aroma. (A) Expression profile of phasiRNA L9_S_361 in fruits of different ripening stages (left) and cleavage validation of transcript Solyc07g006680 coding for an Acyl transferase enzyme (right). (B) Expression profile of phasiRNA L18_AS_1608 in fruits of different ripening stages (left) and cleavage validation of transcript Solyc07g006680 coding for Alcohol dehydrogenase 12 enzyme (right). The red dot indicates the normalized reads that match the predicted cleavage positions mediated by these phasiRNAs at each corresponding transcript in degradome libraries. Peak category and *p-value* according to CleaveLand are indicated above T-plots. IG: immature green, Br: breaker, Pk: pink, RR: red ripe. Different letters indicate a significant difference (adjusted p-value < 0.05) according to edgeR analysis.

The flavor and aroma of different fruits are conferred by a range of different volatile components, synthesized from several different biochemical pathways which include the action of alcohol dehydrogenase (ADH) acyl-transferase (AAT) enzymes (Speirs *et al*. 1998; Lewis *et al*. 2021). These enzymes are involved in the production of alcohols and aldehydes, compounds linked to fruit aromas (Singh *et al*. 2018; Li *et al*. 2021).

#### 3.2.3 PhasiRNAs regulation of fruit softening

Four of the phasiRNAs potentially involved in fruit development and ripening are generated from loci *PHAS3, PHAS14, PHAS19* and *PHAS20,* for which target validation indicates they regulate genes involved in cell wall disassembly and remodeling: L3_AS_452/Xyloglucan endotransglucosylase; L14_S_335/Beta 1,4-xylosidase; and L19_S_2439 and L20_S_988 regulating Polygalacturonases (Table 2).

The expression profile of L3_AS_452 shows a higher accumulation in IG and Br stages, decreases afterwards, showing null expression in RR fruits (Fig. 5A, left), while L19_S_2439 expression follows a similar pattern during ripening, with high accumulation in IG stage only and lower levels in Br, Pk ad RR (Fig. 5B, left). Cleavage site signatures for L3_AS_452 are clearly above background levels (Fig. 5A, T-plot), validating this cleavage event, as well as L19_S_2439 signatures in the transcript coding for a Polygalacturonase enzyme which are especially highly enriched in the predicted cleavage site (Fig. 5B, T-plot).

**Figure 5:**
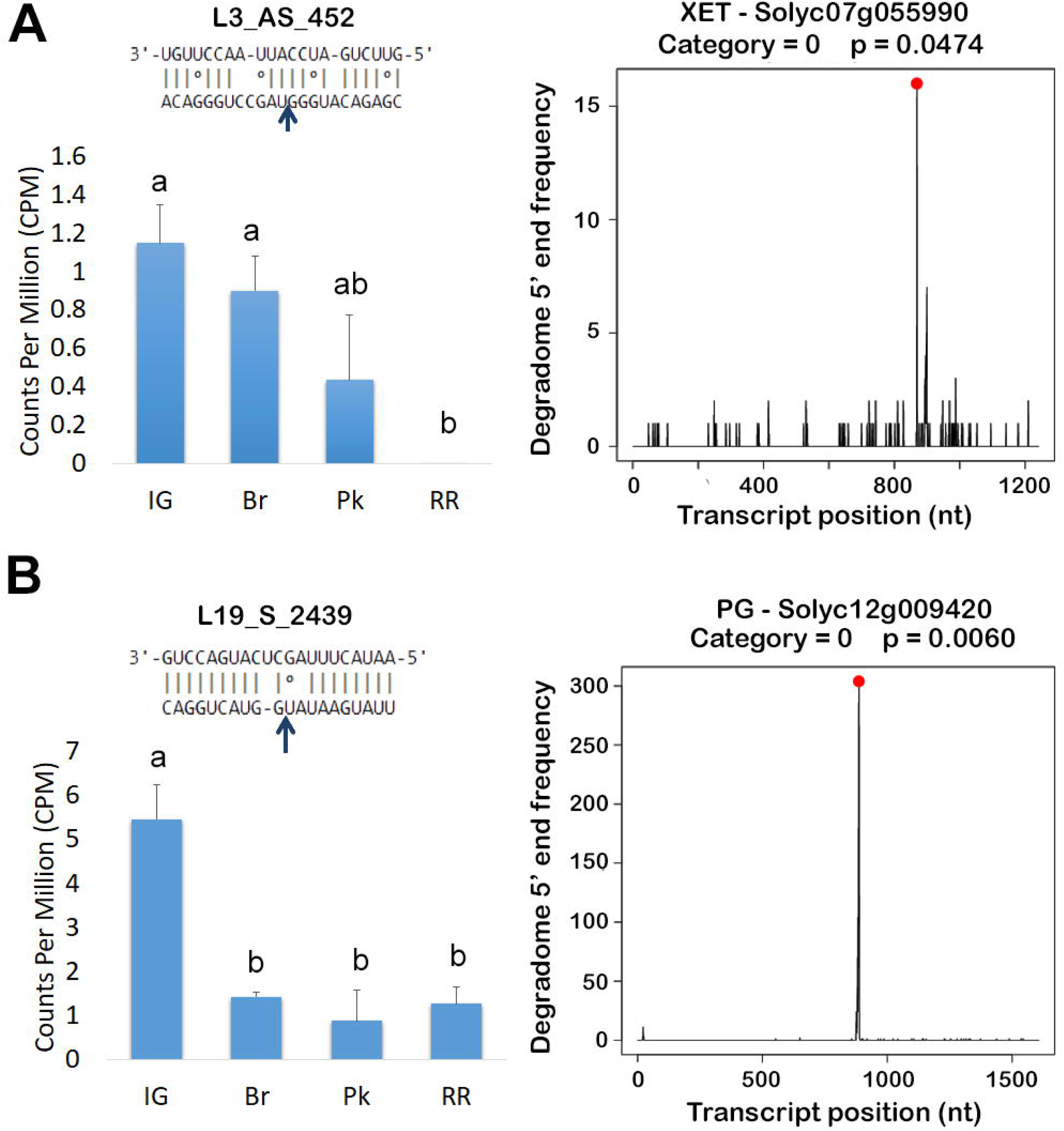
Fruit softening. (A) Expression profile of phasiRNA L3_AS_452 in fruits of different ripening stages (left) and cleavage validation of transcript Solyc07g055990 coding for a Xyloglucan endotransglucosylase enzyme (right). (B) Expression profile of phasiRNA L19_S_2439 in fruits of different ripening stages (left) and cleavage validation of transcript Solyc12g009420 coding for a Polygalacturonase enzyme (right). The red dot indicates the normalized reads that match the predicted cleavage positions mediated by these phasiRNAs at each corresponding transcript in degradome libraries. Peak category and *p-value* according to CleaveLand are indicated above T-plots. IG: immature green, Br: breaker, Pk: pink, RR: red ripe. Different letters indicate a significant difference (adjusted p-value < 0.05) according to edgeR analysis.

These regulatory modules are also attractive for their potential role in the modulation of quality aspects of fruit development.

#### 3.2.4 PhasiRNAs regulating ripening-related transcription factors

Target gene validation analysis identified Solyc03g114840.3.1 as a target gene for phasiRNAs L18_AS_270, derived from locus *PHAS18* (Table 2), and transcript Solyc11g013470.1.1 was also validated as a target for L25_S_1866, derived from locus PHAS25 (Table 2). These transcripts code for transcription factors from SlMADS1 and SlARF17, have been shown to participate in the regulation of several aspects of fruit ripening (Dong *et al*. 2013; Li *et al*. 2023; Mekontso *et al*. 2023).

Analysis of expression levels during fruit ripening shows that phasiRNA L18_AS_270 accumulates in the different ripening stages at levels approximately constant (Fig. 6A, left). The T-plot obtained for this transcript event shows a high peak of signatures that coincides with the L18_AS_270-mediated cleavage site (Fig. 6A, T-plot). Similarly, L25_S_1866 also accumulates at roughly constant levels during ripening, (Fig. 6B), with a validation for the cleavage event on Solyc11g013470.1.1 transcript showing a clear, isolated peak with an elevated number of signatures supporting the regulation mediated by L25_S_1866 of SlARF17 expression. (Fig. 6B, T-plot).

**Figure 6:**
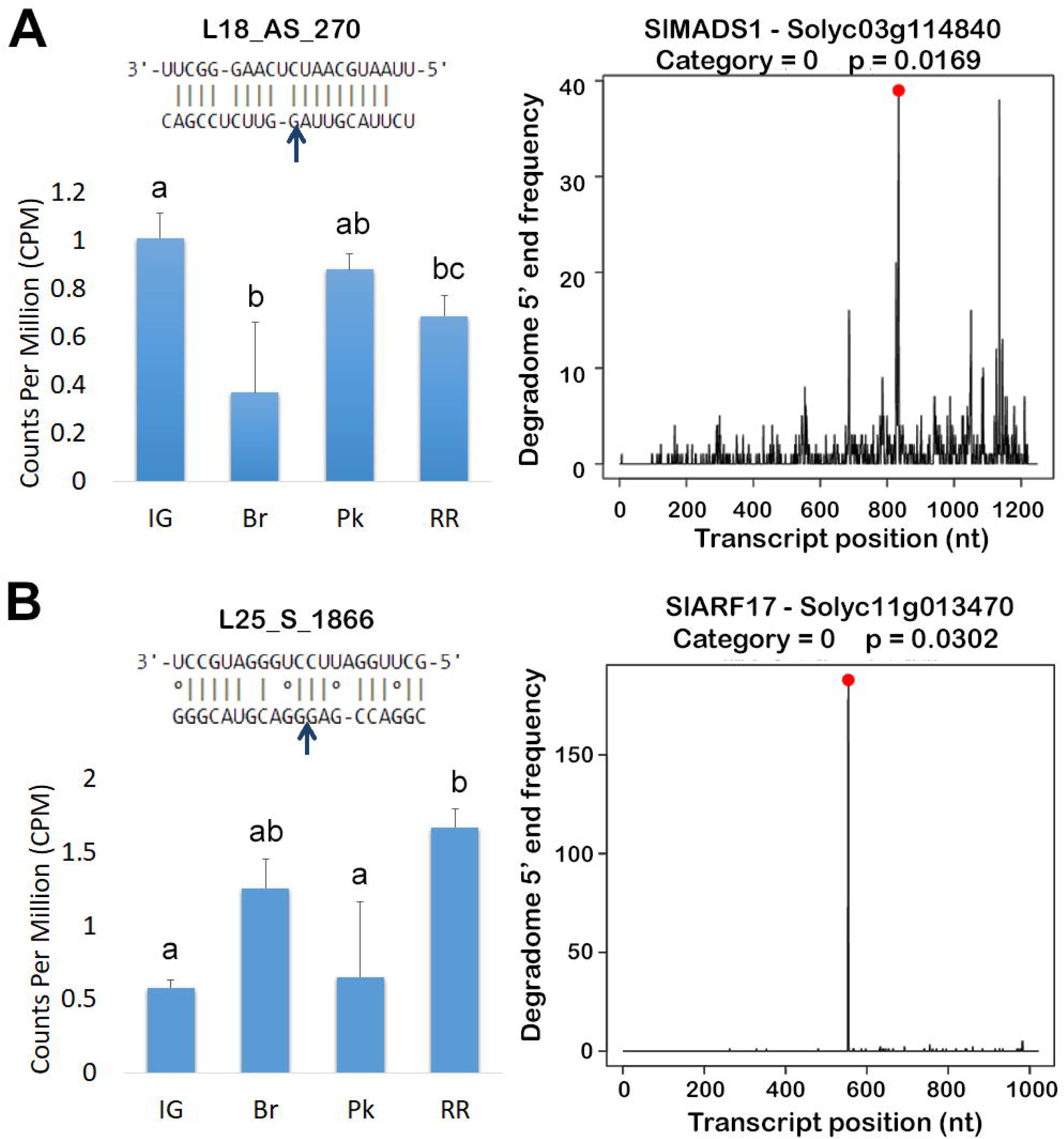
Fruit ripening. (A) Expression profile of phasiRNA L18_AS_270in fruits of different ripening stages (left) and cleavage validation of transcript Solyc03g114840 coding for SlMADS1 transcription factor (right). (B) Expression profile of phasiRNA L25_S_1866 in fruits of different ripening stages (left) and cleavage validation of transcript Solyc11g013470 coding for a SlARF17 transcription factor (right). The red dot indicates the normalized reads that match the predicted cleavage positions mediated by these phasiRNAs at each corresponding transcript in degradome libraries. Peak category and *p-value* according to CleaveLand are indicated above T-plots. IG: immature green, Br: breaker, Pk: pink, RR: red ripe. Different letters indicate a significant difference (adjusted p-value < 0.05) according to edgeR analysis.

### 3.3 Correlation between expression of phasiRNAs and their validated targets

In order to understand the regulatory impact of phasiRNAs on the expression of the validated target transcripts, a comparison of the expression profiles of phasiRNAs and their targets were performed for those targets for which we expect a clear tendency of up- or down- regulation during the ripening process based on their known biological functions and the accumulation patterns of the regulatory phasiRNAs (Figs. 4-6). We analyzed gene expression on IG and RR fruits for targets related to fruit flavor/aroma: ADH - Solyc07g021040.1.1 and AT - Solyc07g006680.1.1, and for those involved in fruit softening: XET - Solyc07g055990.3.1 and PG -Solyc08g082170.4.1 (Fig. 7).

**Figure 7:**
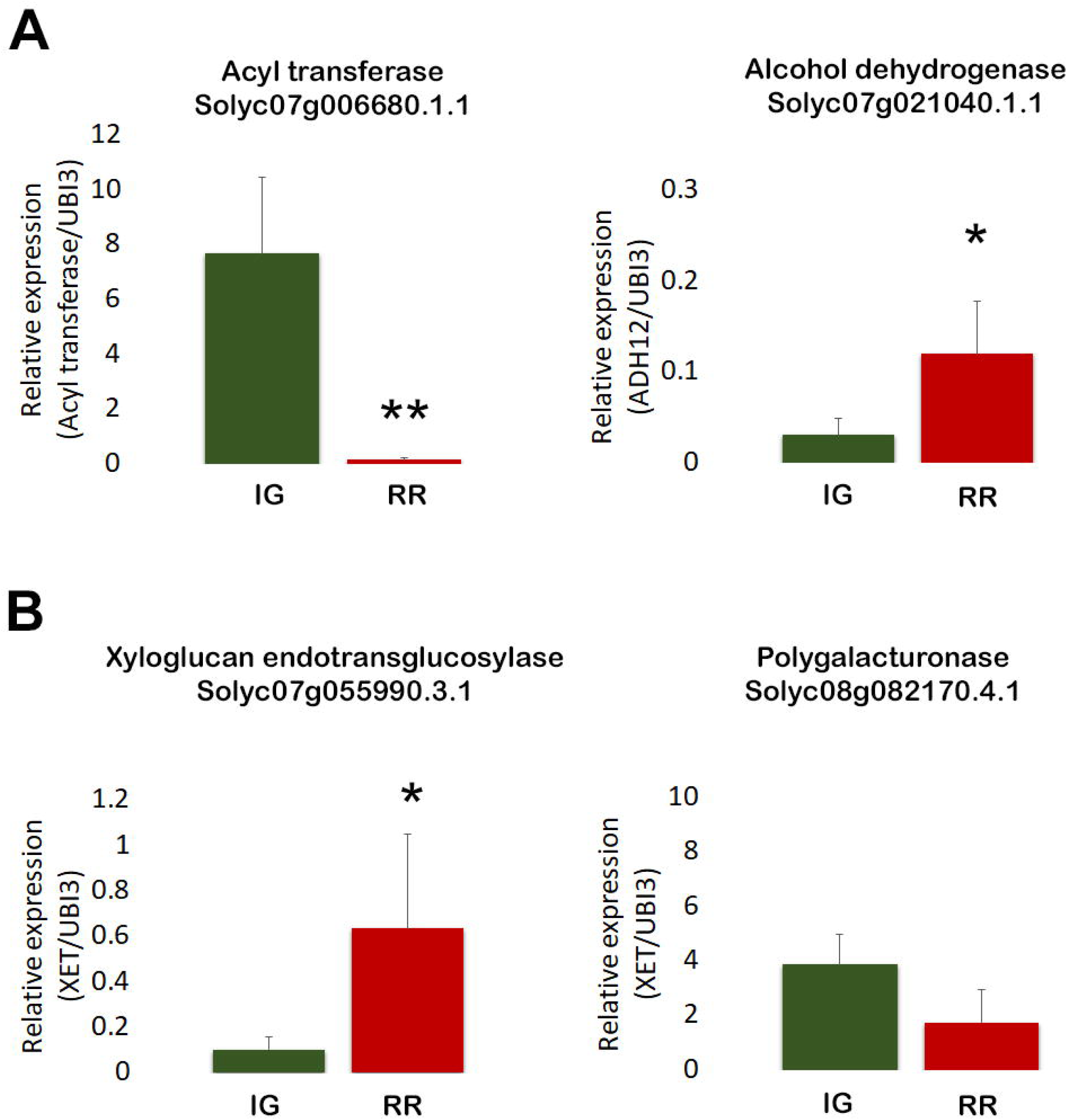
Relative expression of (A) AT and ADH12 genes and (B) XET and PG genes in IG and RR fruits analyzed by RT□ qPCR, using expression of SlUBI3 as internal control. Values represented are mean ± SE (n = 3; Student’s t test, ** = p < 0.001; * = p < 0.05).

As expected, the regulatory modules AT/L9_S_361 and ADH/L18_AS_1608, exhibit an inverse expression pattern to their targets during ripening in the ripening stages analyzed (Fig. 7A). L9_S_361 increases expression between IG and RR stages (Fig. 4A), whereas the AT transcript Solyc07g006680.1.1 decreases its accumulation between these two stages (Fig. 7A). The opposite is observed for the ADH enzyme transcript Solyc07g021040.1.1, which increases its expression from IG to RR (Fig. 7A), with the opposite tendency observed for its regulatory phasiRNA L18_AS_1608 (Fig. 4B).

Similarly, for L3_AS_452/XET, expression of the phasiRNA was high in IG and null in RR fruits (Fig. 5A), while the XET transcript Solyc07g055990.3.1 showed the opposite pattern, increasing its expression from IG to RR stages (Fig. 7B). However, the regulatory module L19_S_2439/PG did not show the expected inverse expression pattern between the phasiRNA and its target PG transcript Solyc08g082170.4.1. For L19_S_2439 a decrease in expression between IG and RR stages was observed (Fig. 5B), while no changes in expression were detected for PG transcript (Fig. 7B).

## 4. Discussion

Previous studies highlight the role of phasiRNAs in the regulation of the expression of defense proteins against biotic stress in various species (Fei *et al*. 2013b; Wu *et al*. 2017) and a study demonstrates the relationship between these small RNAs and abiotic stress, specifically drought resistance, in *Populus trichocarpa* (Shuai *et al*. 2016). PhasiRNAs have also been detected in floral tissues and related to reproduction in several species (Kakrana *et al*. 2018; Xia *et al*. 2019; Liu *et al*. 2020). Also, previous work that analyzed samples from tomato leaves detected 91 *PHAS* loci out of which 55 phasiRNAs generated from these *loci* increased expression in the presence of the fungus *Botrytis cinerea*, and a total of 137 phasiRNAs were also identified in tomato leaves, most of which are produced from defense genes of the NBS-LRR type and whose expression is modified in response to abscisic acid (Wu *et al*. 2017; Luan *et al*. 2020). However, no reports were found that analyze phasiRNAs in fruits and link them to their ripening or the development of organoleptic properties and fruit quality traits.

In this work, 25 *PHAS* loci generating phasiRNAs expressed during tomato fruit ripening were identifies for the first time and using degradome analysis for target validation, we identified several regulatory modules that could have a role in the regulation of fruit ripening and quality. These include genes related to fruit brightness, color, texture and aroma and flavor. The results presented here allow for further exploration and possibly the development of biotechnological tools to improve organoleptic properties in tomato fruits.

The location of 17 of these loci matches protein-coding genes annotated in the tomato genome, and similar to *PHAS* loci detected in tomato leaves, they are associated to proteins involved in plant’s defense and were not further analyzed.

Interestingly, our identification of miRNA triggers for the phasiRNAs derived from these loci, identified only miR482 to trigger phasiRNAs from *PHAS2, PHAS4* and *PHAS22*. However, triggers for phasiRNA production have not been identified in many cases and is possible that additional mechanisms exist. Since Dicer-Like enzymes are capable of generating small RNAs from several types of dsRNA (Henderson *et al*. 2006; Margis *et al*. 2006; Fukudome and Fukuhara 2017), one possibility that arises from exploring the genomic locations where some of these *PHAS* loci are localized, is that overlapping antisense transcripts are simultaneously expressed (e.g. *PHAS3* and *PHAS19*, Suppl. File 3) and a resulting dsRNA can be processed by DCL enzymes to generate 21-nt phasiRNAs. Alternatively, some of these *PHAS* loci are located in regions where lncRNAs are annotated (see Table 1), which could possibly fold to adopt secondary structures with dsRNA regions, which could also be processed into phasiRNAs by DCL enzymes.

A total of 186 target genes regulated by phasiRNAs derived from these loci were validated, among which we selected 13 that would have a direct relationship with the development and acquisition of organoleptic properties or quality traits in tomato fruits (Table 2).

Among the target genes identified, two phasiRNAs were selected for being validated as regulators of an enzyme of the GDSL lipase family, directly related to the formation of the fruit cuticle and therefore to protective properties of fruit integrity and the acquisition of fruit brightness (Petit *et al*. 2014). It should be noted that this group of enzymes has not been studied in detail in tomatoes, so it is interesting to characterize them in the future in relation to their influence on the appearance of the fruit and its regulation by this small RNA.

Related to fruit color, phasiRNA L2_S_505 was selected for its regulation of the transcript coding for Phototropin 1. These proteins are UVA-blue light photoreceptors that detect the direction of light and optimize photosynthesis, but they also play roles in other developmental responses. Phototropin 1 is particularly interesting because tomato mutants in this gene were found to have altered content of carotenoids and lycopene in particular, which is the main compound responsible for red color in tomatoes (Kilambi *et al*. 2021). Hence, this regulatory module also represents an interesting possibility for the fine tuning of fruit color in tomatoes.

The main taste characteristics of fruit are derived from the relative concentrations of organic acids and sugars (Kader 2008; Bauchet *et al*. 2017). Alcohol acetyltransferases are important in volatile ester synthesis and the final step in the pathway, conversion of aldehydes to alcohols, requires ADH activity (Yahyaoui *et al*. 2002; Tieman *et al*. 2012). PhasiRNAs were also found to regulate genes related to the production of volatile compounds, such as ADH enzymes. There are reports linking this protein to aroma production in several species (Echeverrı a *et al*. 2004; Manríquez *et al*. 2006) and in tomatoes, this family of enzymes has been directly related to the taste of fruits (Speirs *et al*. 1998; Yilmaz *et al*. 2002), but there are few molecular studies that attempt to improve this attribute through genetic modifications. Due to the large number of compounds involved in the acquisition of flavor, the modification of only one or a few does not produce substantial changes in the perception of this property, so the use of genetic engineering techniques for breeding, where the expression of a single gene is modified, has not been very effective. or modification of these genes. With this in mind, the use of several phasiRNAs with the ability to regulate the expression of ADH or AT transcripts appears as an interesting possibility for further analysis.

Fruit softening is an irreversible process that is part of the fleshy fruit ripening process and is partially due to cell wall changes catalyzed by wall-modifying enzymes. However, excessive softening reduces shelf life, increases the possibility of physical damage and the subsequent susceptibility to pathogen attack. Also, excessive softening is typically a fruit quality not preferred by consumers (Klee and Giovannoni 2011; Zuo *et al*. 2012; Klee and Tieman 2013). Polygalacturonases are enzymes involved in the disassembly of the polysaccharide network of the cell wall during fruit ripening, thus participating in fruit softening, a key sensory aspect related to fruit quality and consumer acceptance (Jiang *et al*. 2019). These enzymes hydrolyze the glycosidic α-1,4 bonds between galacturonic acid residues, components of the pectin network of the plant cell wall, producing the degradation of the polysaccharide matrix and contributing to fruit softening (Fry 2004; Illera *et al*. 2018). It was also found that in tomatoes with reduced activity of these proteins, post-harvest life improved, as did the resistance of the fruits to transport (Schuch *et al*. 1991). In this study, expression of this gene was not found to present the expected inverse correlation between phasiRNA L19_S_2439 and its target transcript, perhaps because regulation by this phasiRNA is not the main one for this transcript, in spite of validated cleavage event. The expected inverse expression correlation was observed for ’phasiRNA L3_AS_452, which modulates the expression of another cell wall remodeling enzyme, Xyloglucan Endotransglucosylase. These enzymes have been proposed to have a dual role integrating newly secreted xyloglucan chains into an existing wall-bound xyloglucan, or restructuring the existing cell wall material by catalyzing transglucosylation between xyloglucan molecules, for which a decrease in activity during ripening was proposed to contribute to the fruit softening (Saladié *et al*. 2006; Miedes and Lorences 2009; Miedes *et al*. 2010).

Hence, the regulatory modules identified here could also be used to reduce the accumulation of transcripts encoding PG and XET enzymes, allowing for a modulation in the disassembly of the cell wall to generate fruits with longer postharvest life.

Finally, phasiRNAs regulating two transcription factors were validated, SlMADS1 from the MADS-box family and SlARF17, part of the AUXIN RESPONSE FACTORS family of proteins. Even though tomato is a climacteric fruit where most aspects related to ripening are regulated by ethylene, it has also been shown that auxins are involved in the control of sugar metabolism during tomato ripening (Sagar *et al*. 2013; Mekontso *et al*. 2023). In addition, another study showed that ARFs genes (particularly SlARF2) would positively regulate fruit ripening (Hao *et al*. 2015). This family of proteins has already been characterized in tomato (Kumar *et al*. 2011) but there are no studies reporting regulation by phasiRNAs, making the regulation of ARF17 expression mediated by L25_S_1866 interesting for its potential role in flavor determination and to modulate tomato fruit ripening. Similarly, SlMADS1 is one of many transcription factors from the MADS family that was shown to participate in the regulation of several aspects of tomato fruit ripening, (Dong *et al*. 2013; Li *et al*. 2023), for which L18_AS_270 represents another interesting candidate for further testing.

Taking together, the results presented here provide a number of phasiRNA/transcript regulatory modules that are useful as tools for the genetic improvement of attributes of agronomic interest related to the qualitative aspects of tomato fruits and to the organoleptic properties attractive to consumers. These results provide evidence for siRNA-guided post-transcriptional regulation of target genes involved in tomato fruit development, highlighting their dynamic expression patterns and functional significance during ripening.

## Author Contributions

M. D. was responsible for the study conception and design. Material preparation, data collection and analysis were performed by S. S. and A. S. The first draft of the manuscript was written by M. D. and all authors commented on previous versions of the manuscript. All authors read and approved the final manuscript.

## Data Availability

The datasets generated and/or analyzed during the current study are included in the manuscript or as supplemental materials.

## Supplemental material

Supplemental File 1: Libraries used and mapping statistics

Supplemental File 2: PhasiRNA sequences

Supplemental File 3: PHAS loci genomic locations

Supplemental File 4: Validated phasiRNA targets

