## Supplemental File 3 for "Genome-wide analysis of phased small interfering RNAs related to tomato fruit ripening and quality"

*PHAS1*: SL4.0 ch01:4415780-4416701

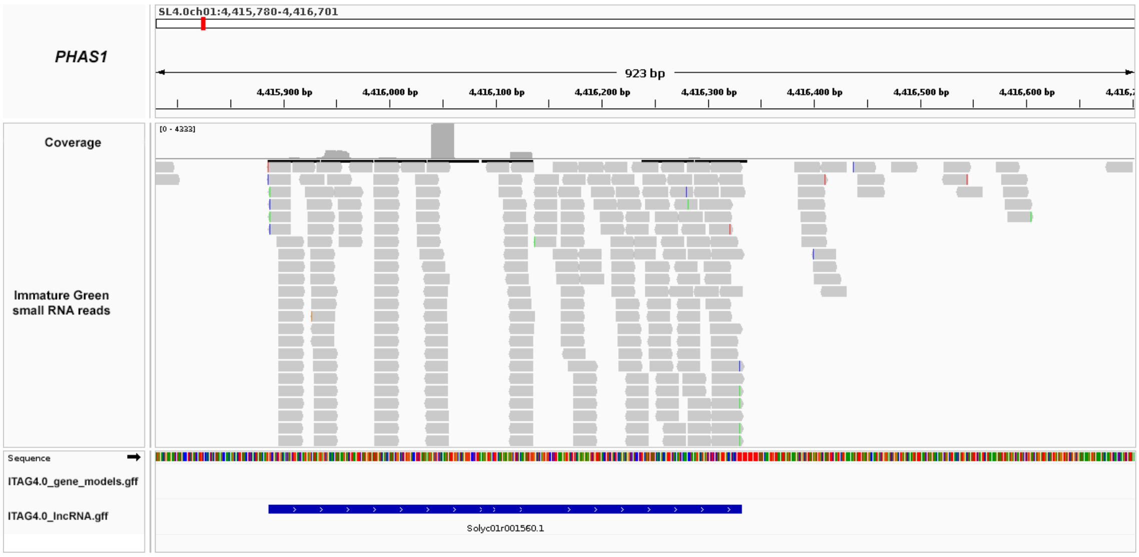


*PHAS2*: SL4.0 ch02:29160966-29163434

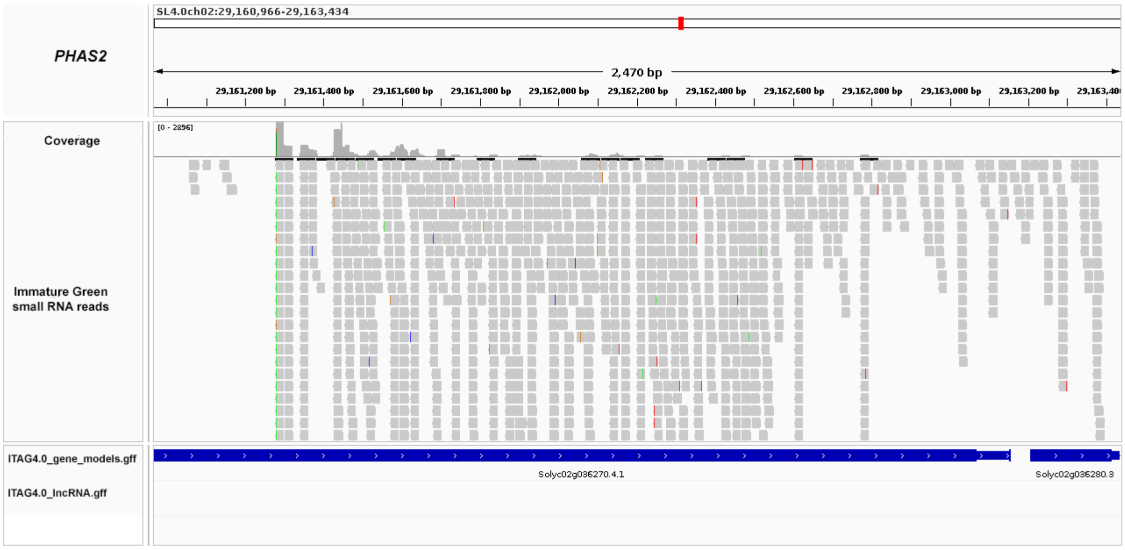


*PHAS3*: SL4.0 ch04:373826-376280

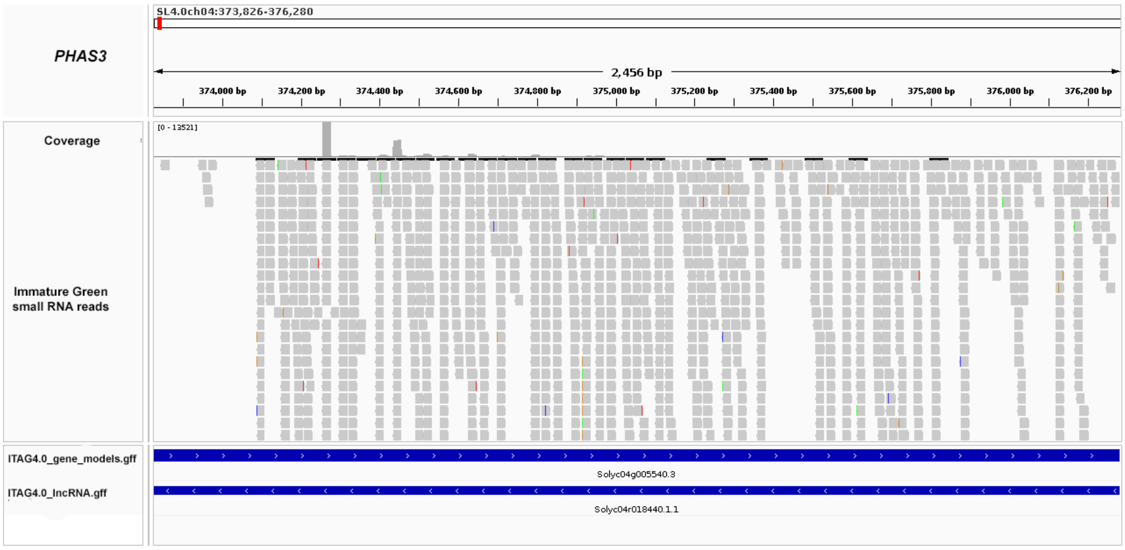

*PHAS4*: SL4.0 ch05:2549891-2552244

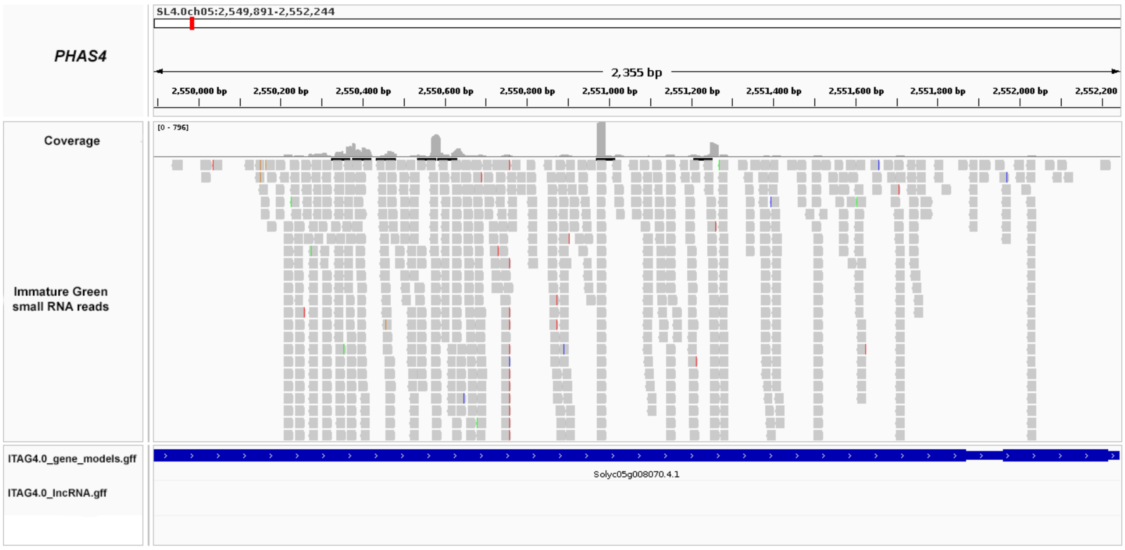


*PHAS5*: SL4.0 ch05:3872005-3874078

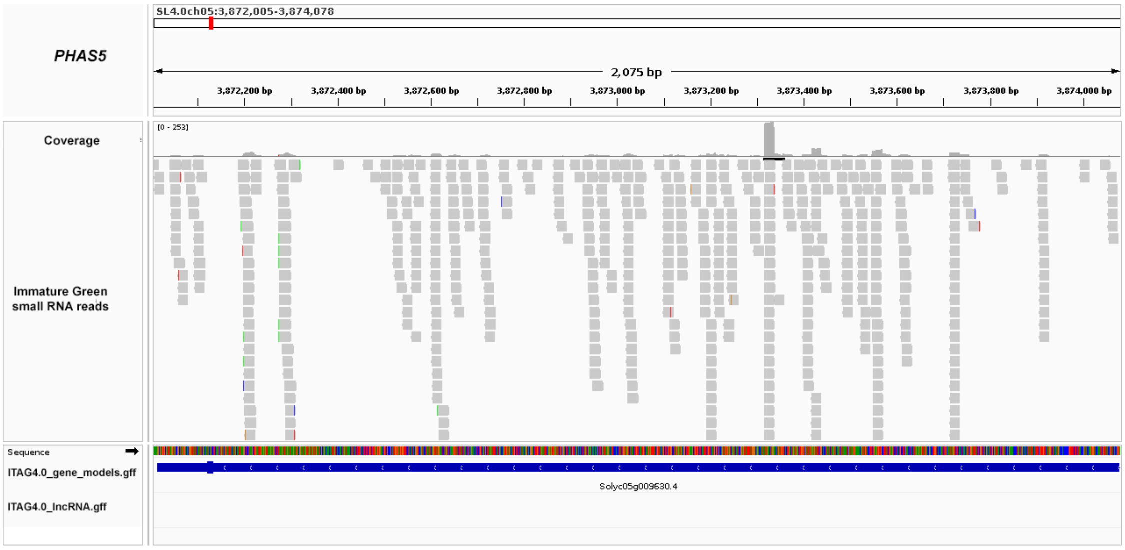


*PHAS6*: SL4.0 ch06:42183189-42183665

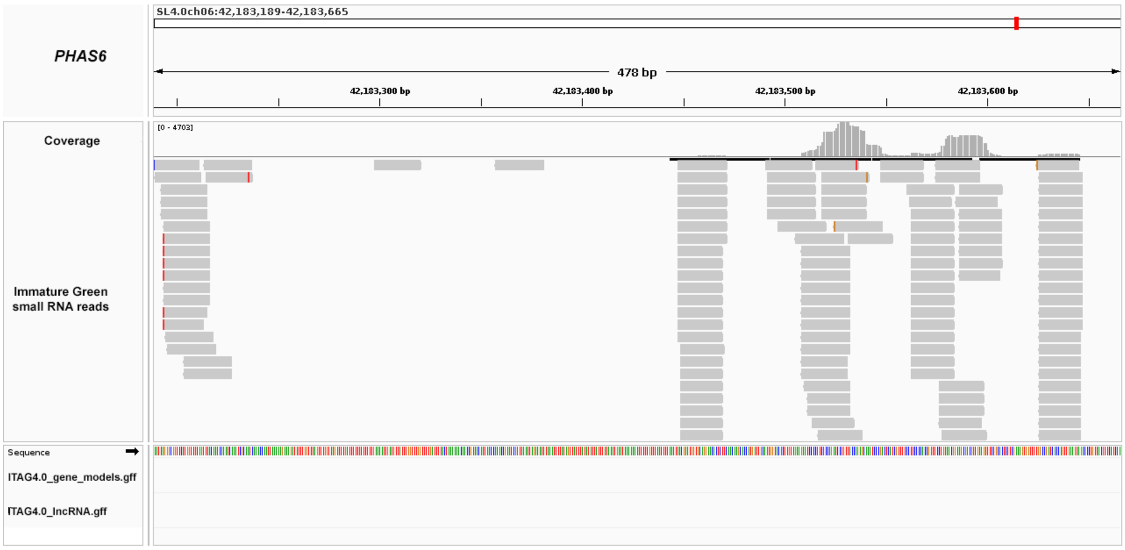

*PHAS7*: SL4.0 ch06:42207523-42208019

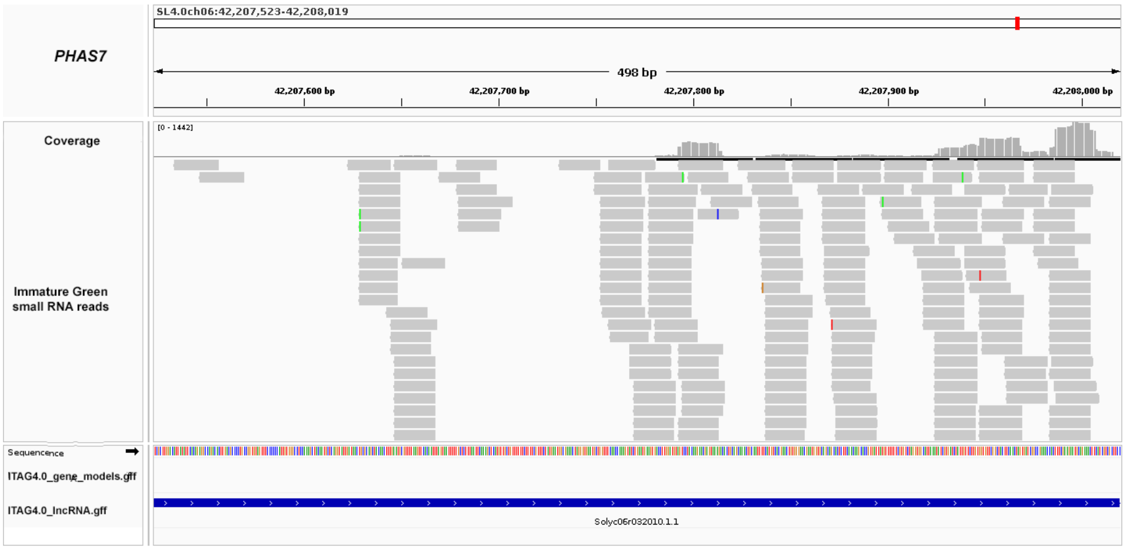


*PHAS8*: SL4.0 ch06:437853-438581

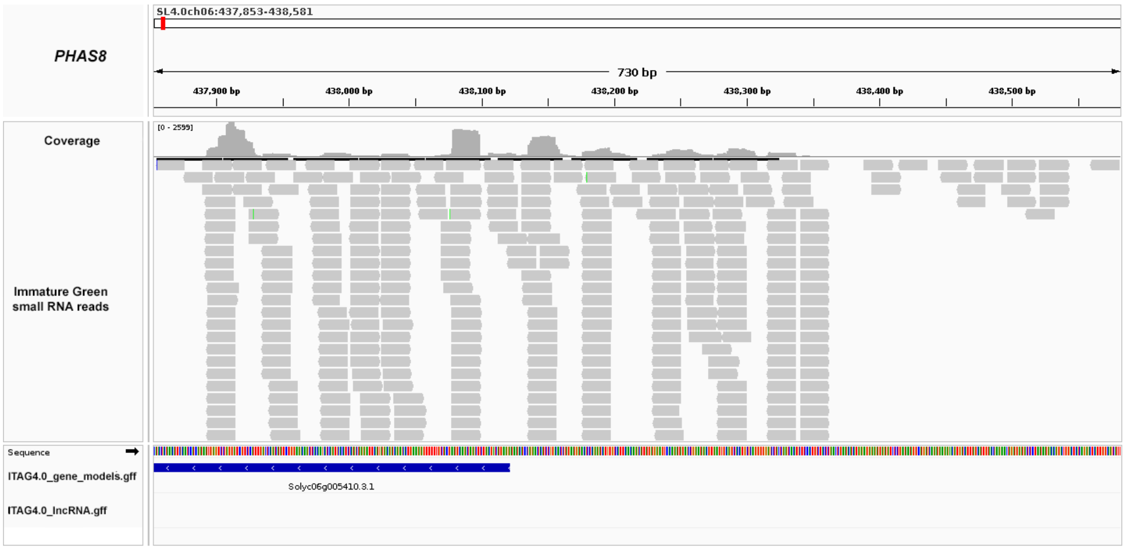


*PHAS9*: SL4.0 ch06:439060-439697

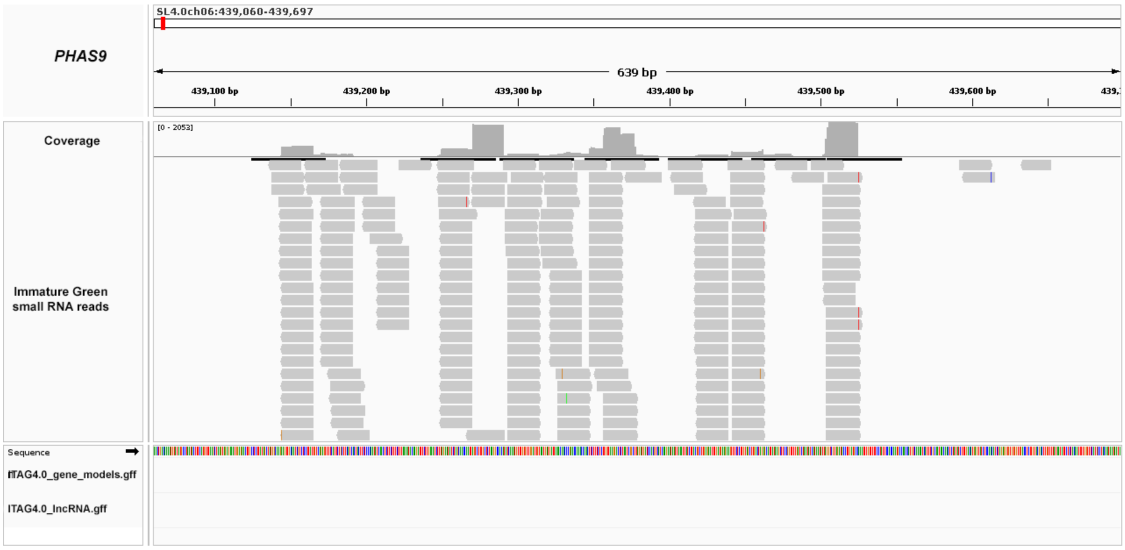

*PHAS10*: SL4.0 ch06:704215-704566

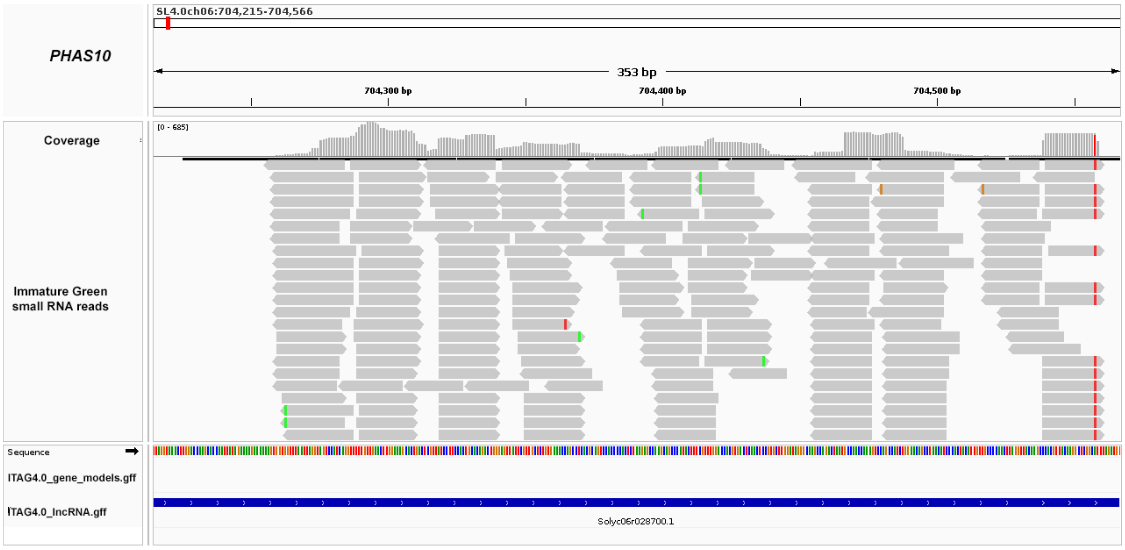


*PHAS11*: SL4.0 ch07:4205316-4207077

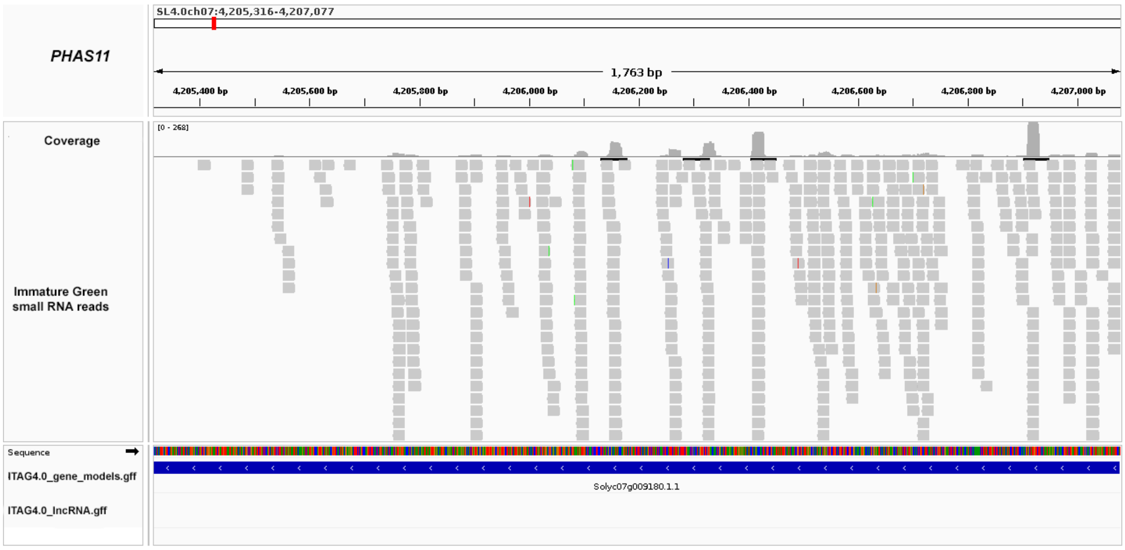


*PHAS12*: SL4.0 ch08:13837317-13838009

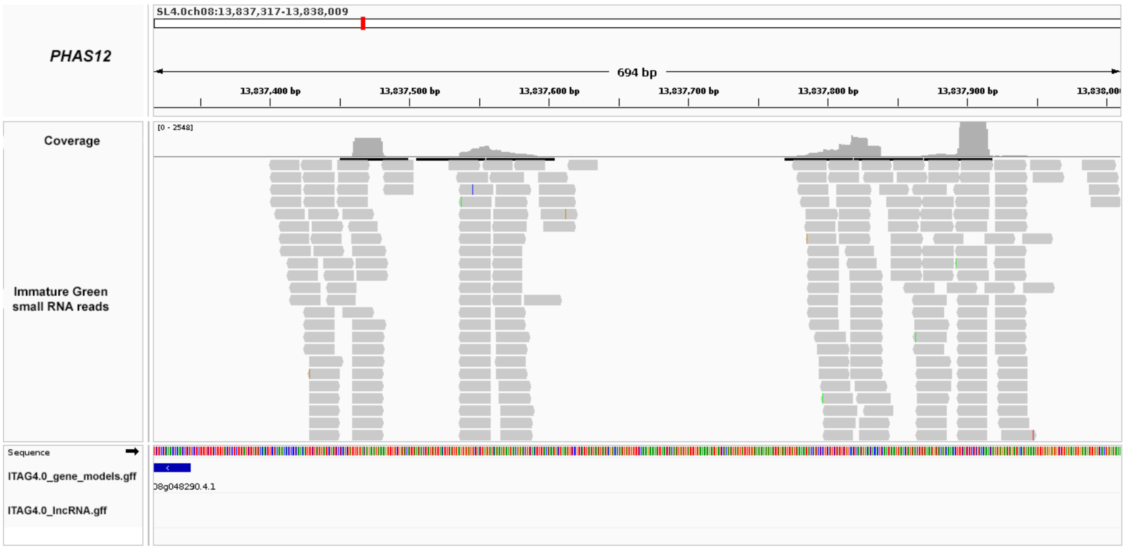

*PHAS13*: SL4.0 ch08:57063495-57067399

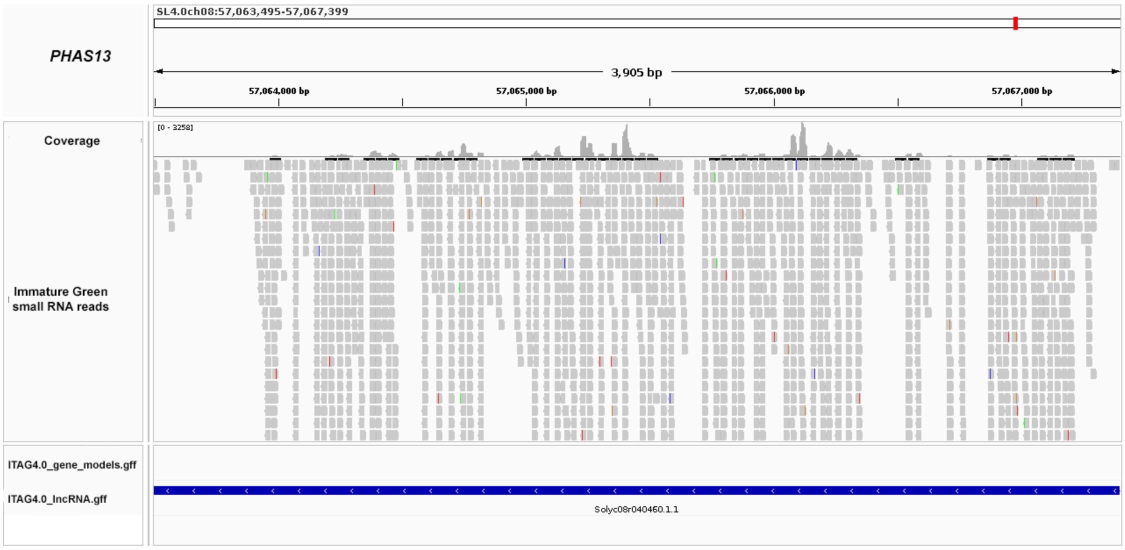


*PHAS14*: SL4.0 ch08:59208402-59210241

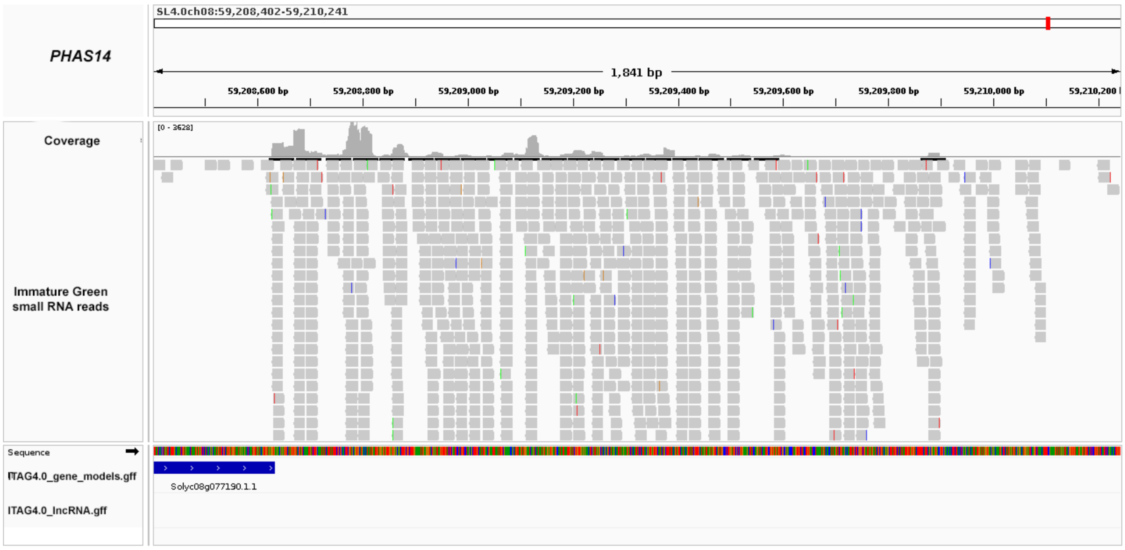


*PHAS15*: SL4.0 ch09:13657071-13658918

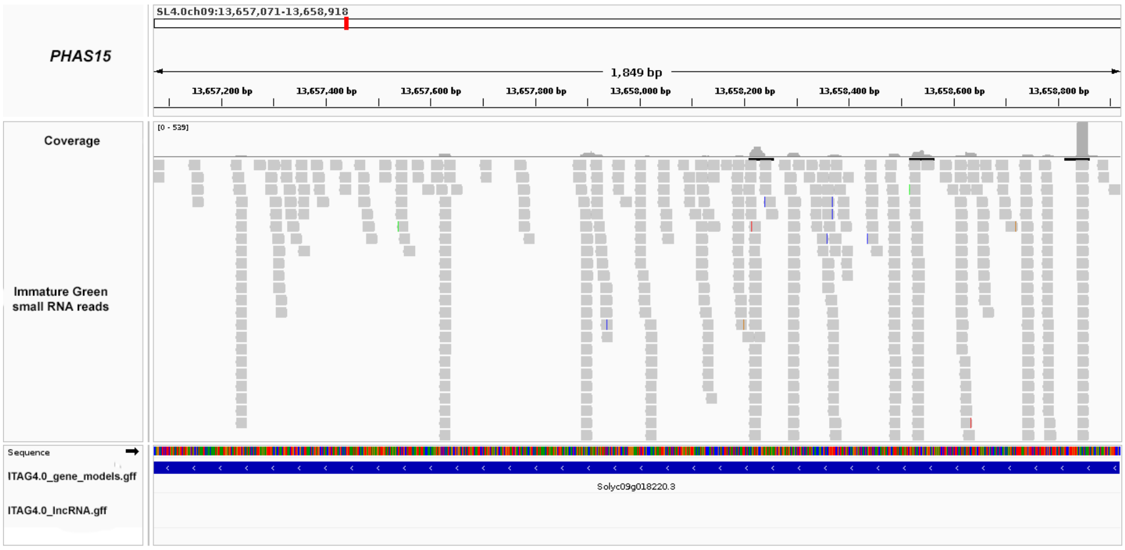

*PHAS16*: SL4.0 ch09:57495810-57497595

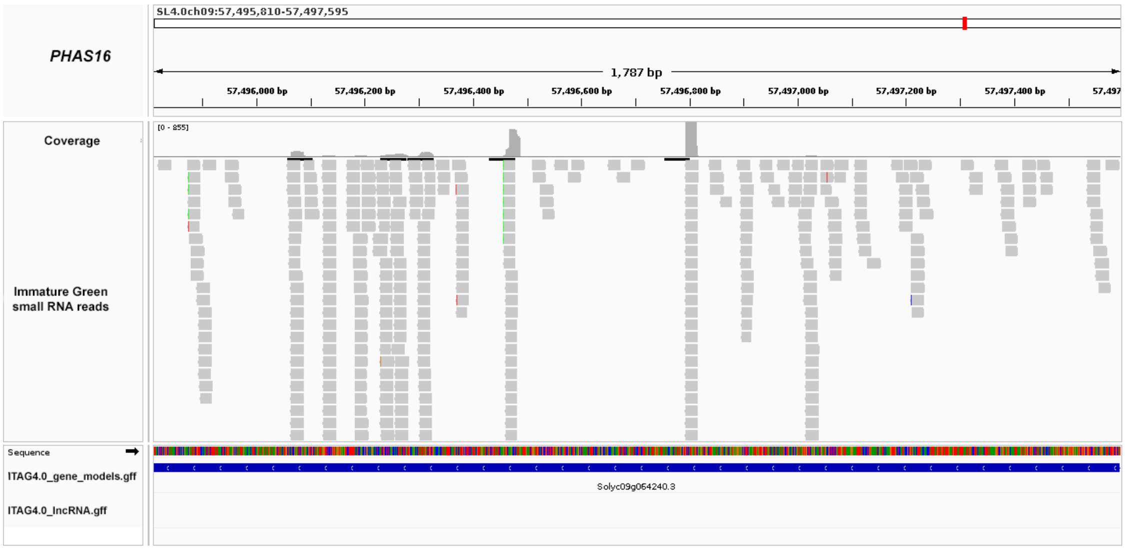


*PHAS17*: SL4.0 ch10:58274198-58274971

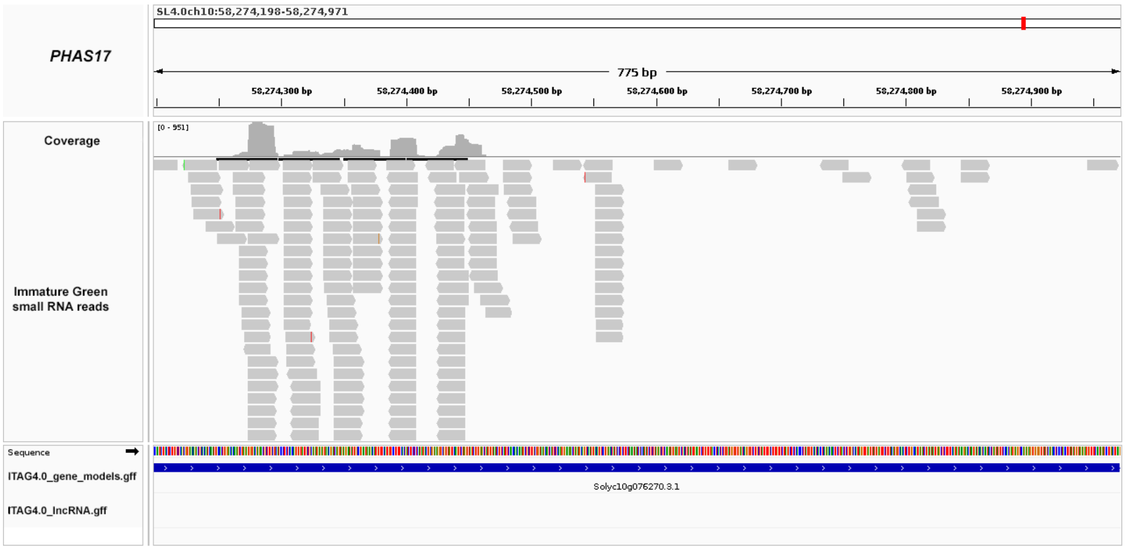


*PHAS18*: SL4.0 ch11:2733015-2740117

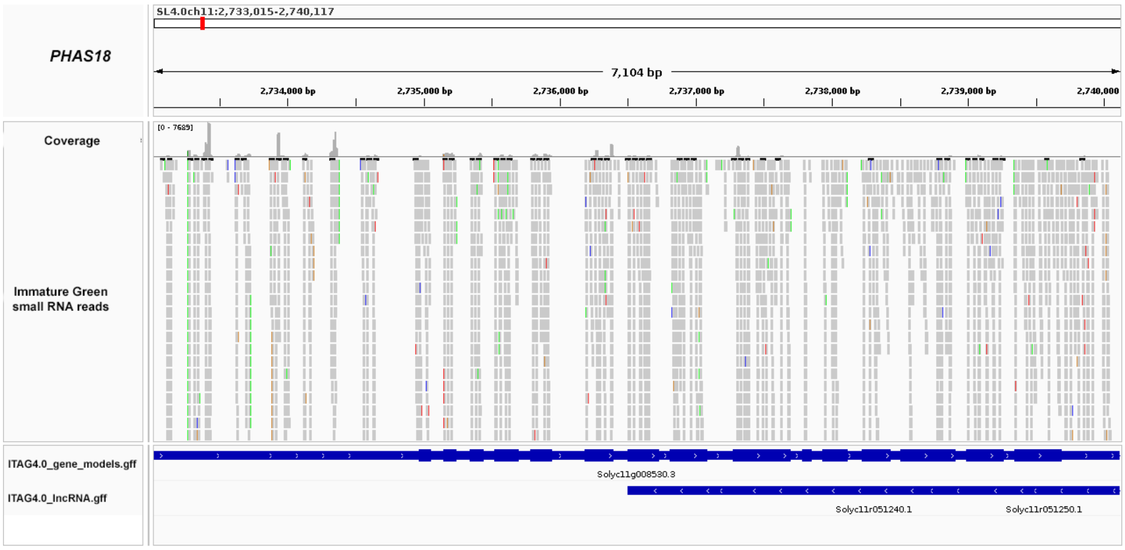

*PHAS19*: SL4.0 ch11:2747539-2754315

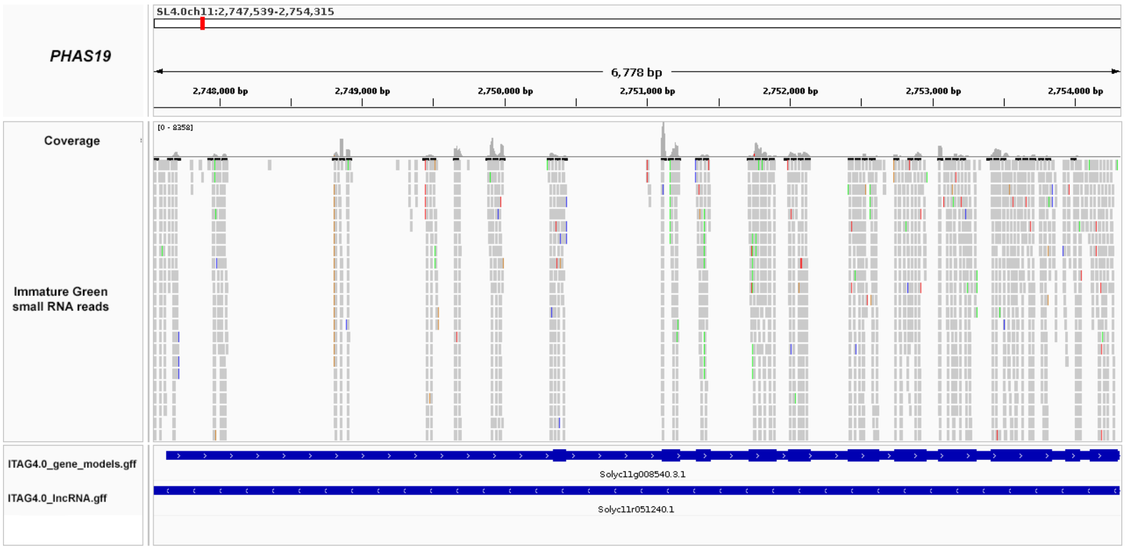


*PHAS20*: SL4.0 ch11:49432234-49436209

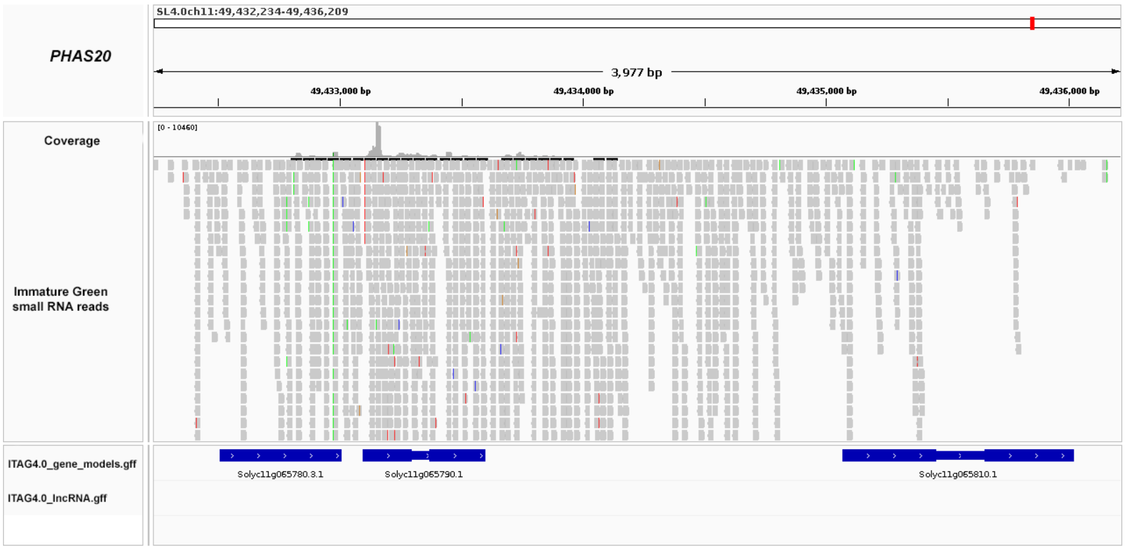


*PHAS21*: SL4.0 ch11:52640011-52642954

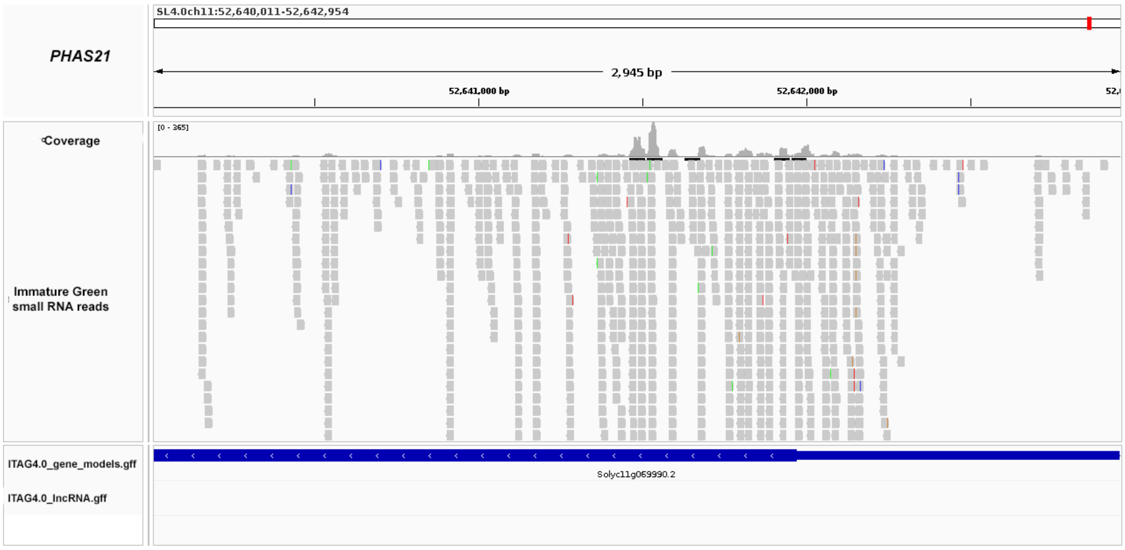

*PHAS22*: SL4.0 ch11:7217527-7218901

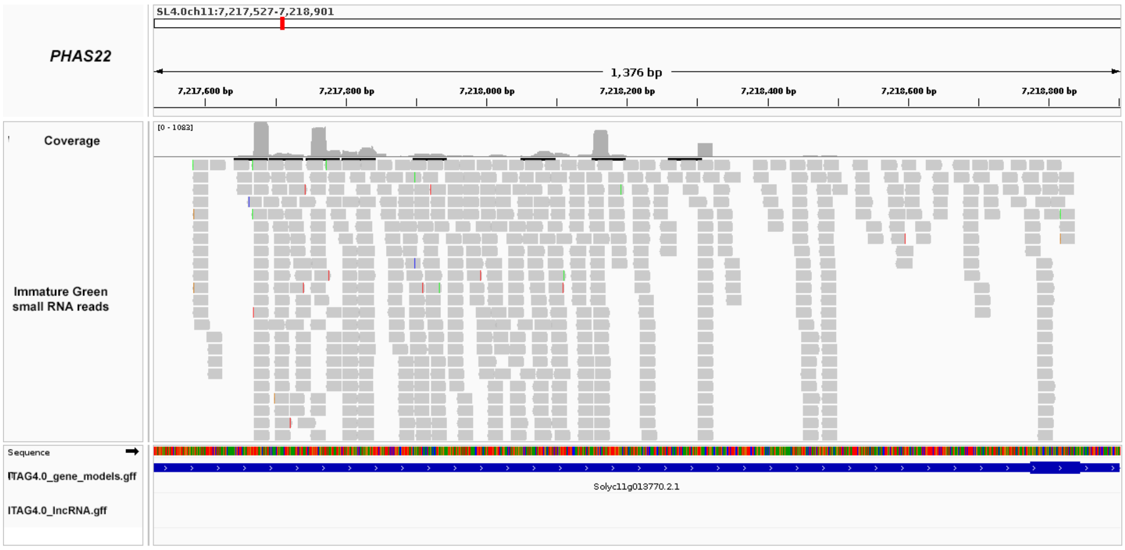


*PHAS23*: SL4.0 ch12:2829642-2831621

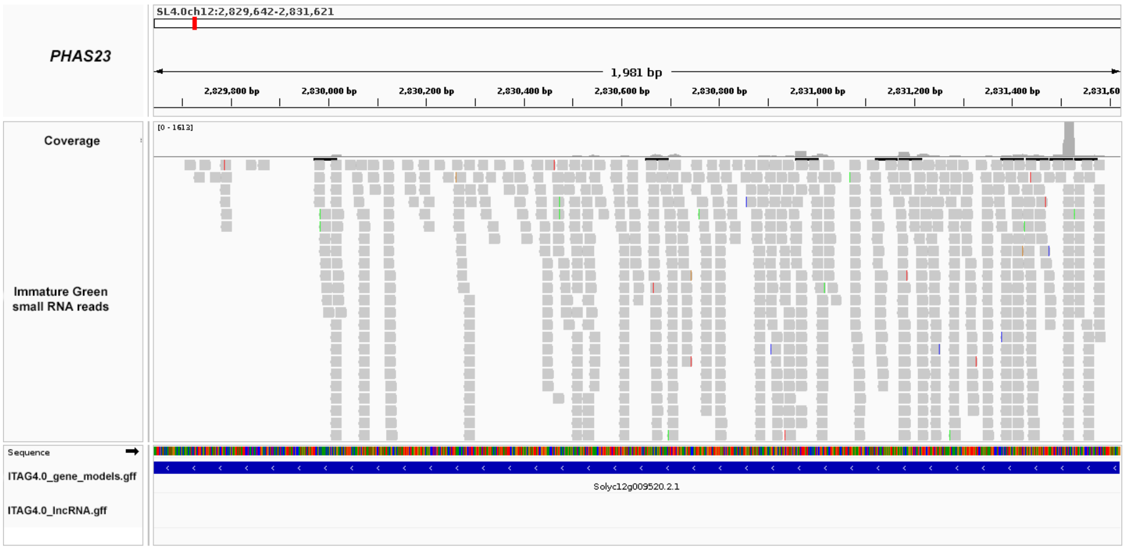


*PHAS24*: SL4.0 ch12:641458-644168

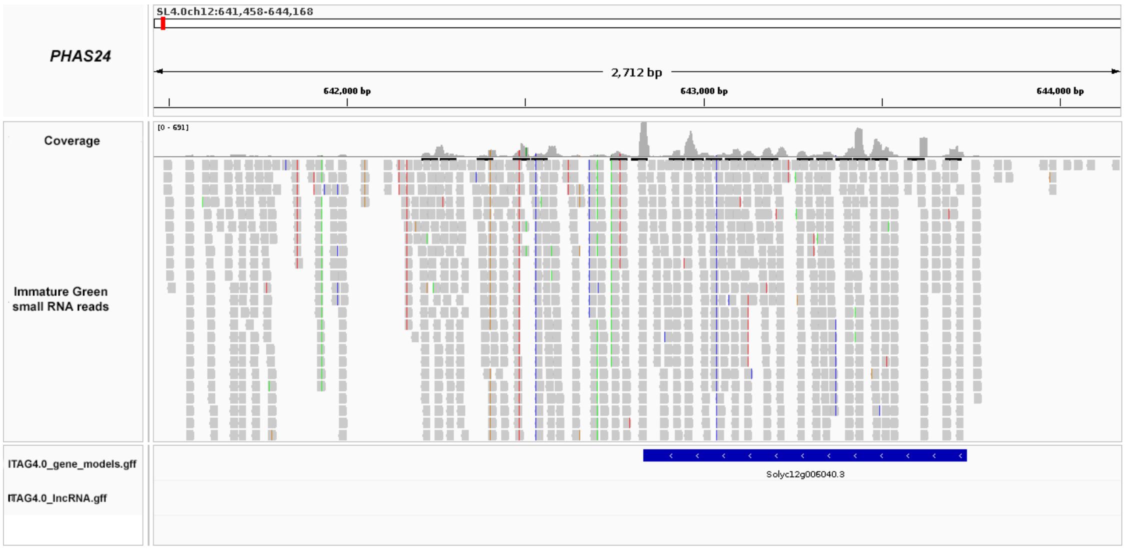

*PHAS25*: SL4.0 ch12:6447225-6449223

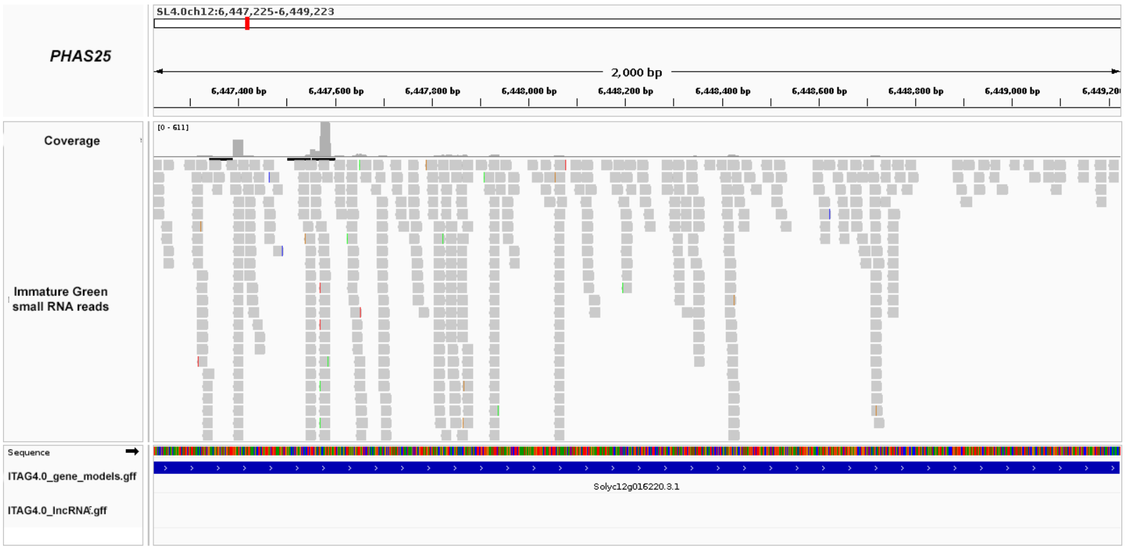
